# Epilepsy-associated *SCN2A* (Na_V_1.2) Variants Exhibit Diverse and Complex Functional Properties

**DOI:** 10.1101/2023.02.23.529757

**Authors:** Christopher H. Thompson, Franck Potet, Tatiana V. Abramova, Jean-Marc DeKeyser, Nora F. Ghabra, Carlos G. Vanoye, John Millichap, Alfred L. George

**Affiliations:** Department of Pharmacology, Feinberg School of Medicine, Northwestern University, Chicago, IL 60611 USA; Department of Neurology, Feinberg School of Medicine, Northwestern University, Chicago, IL 60611 USA

**Keywords:** sodium channel, Na_V_1.2, SCN2A, epilepsy, electrophysiology

## Abstract

Pathogenic variants in neuronal voltage-gated sodium (Na_V_) channel genes including *SCN2A*, which encodes Na_V_1.2, are frequently discovered in neurodevelopmental disorders with and without epilepsy. *SCN2A* is also a high confidence risk gene for autism spectrum disorder (ASD) and nonsyndromic intellectual disability (ID). Previous work to determine the functional consequences of *SCN2A* variants yielded a paradigm in which predominantly gain-of-function (GoF) variants cause epilepsy whereas loss-of-function (LoF) variants are associated with ASD and ID. However, this framework is based on a limited number of functional studies conducted under heterogenous experimental conditions whereas most disease-associated *SCN2A* variants have not been functionally annotated. We determined the functional properties of more than 30 *SCN2A* variants using automated patch clamp recording to assess the analytical validity of this approach and to examine whether a binary classification of variant dysfunction is evident in a larger cohort studied under uniform conditions. We studied 28 disease-associated variants and 4 common population variants using two distinct alternatively spliced forms of Na_V_1.2 that were heterologously expressed in HEK293T cells. Multiple biophysical parameters were assessed on 5,858 individual cells. We found that automated patch clamp recording provided a valid high throughput method to ascertain detailed functional properties of Na_V_1.2 variants with concordant findings for a subset of variants that were previously studied using manual patch clamp. Additionally, many epilepsy-associated variants in our study exhibited complex patterns of gain- and loss-of-function properties that are difficult to classify overall by a simple binary scheme. The higher throughput achievable with automated patch clamp enables study of a larger number of variants, greater standardization of recording conditions, freedom from operator bias, and enhanced experimental rigor valuable for accurate assessment of Na_V_ channel variant dysfunction. Together, this approach will enhance our ability to discern relationships between variant channel dysfunction and neurodevelopmental disorders.

## INTRODUCTION

Epilepsy and neurodevelopmental disorders (NDD) with clinical onset during infancy and childhood are often attributed to monogenic etiologies (1-3). Among the dozens of individual genes associated with these conditions are several that encode voltage-gated ion channels including sodium and potassium channels expressed throughout the nervous system. Defining the functional consequences of ion channel variants in this disease context can inform a mechanistic framework that advances understanding of pathophysiology and helps guide conceptualization of targeted therapeutic approaches (4).

*SCN2A* encodes a voltage-gated sodium channel (Na_V_1.2) that is expressed widely in developing and mature brain. Pathogenic *SCN2A* variants are associated with childhood-onset epilepsy of varying severity as well as autism spectrum disorder (ASD) with or without accompanying seizures and nonsyndromic intellectual disability (ID) (5-8). The type of variant in these conditions differs to some extent with truncating variants (premature stop codons, frameshifts) being most prevalent in ASD/ID whereas missense variants being more common in persons with epilepsy. Because truncating *SCN2A* variants likely produce nonfunctional Na_V_1.2 channels, and because earlier studies of missense variants associated with epilepsy demonstrated gain-of-function (GoF), a general genotype-phenotype correlation emerged (9). Specifically, evidence from *in vitro* functional evaluation of *SCN2A* variants supported the hypothesis that loss-of-function (LoF) variants were the drivers of ASD/ID, whereas GoF variants caused early-onset epilepsy (10). This genotype-phenotype correlation for *SCN2A* provided a framework to predict which individuals might respond best to sodium channel blocking anti-convulsant drugs. Seizures may also occur in individuals with *SCN2A* LoF variants and evidence from mouse models suggests that maladaptive changes in potassium channel expression in the brain may offer an explanation (11, 12).

Distinguishing GoF and LoF based on *in vitro* studies of recombinant Na_V_1.2 channels can be challenging because of the myriad of functional properties that can be assessed. This is particularly the case with missense variants of uncertain significance. Variants that fail to generate measurable sodium current are easy to categorize as complete LoF. Variants that affect the time course or voltage-dependence of channel gating may also be simple to characterize as GoF in some cases. For example, a depolarizing shift in the voltage-dependence of inactivation or a hyperpolarizing shift in the voltage-dependence of activation will promote greater open probability of sodium channels. Abnormal inactivation (e.g., slower time course, enhanced persistent current) is also recognized as a GoF property of variant sodium channels. When these dysfunctional properties occur in isolation, it is straightforward to assign gain or loss of function to a variant. However, more complex profiles of gain and loss of individual functional properties are more difficult to categorize.

In this study, we determined the functional properties of several *SCN2A* variants mostly associated with neonatal onset epilepsy using automated patch clamp recording, which enables higher throughput than traditional manual patch clamp. We validated this approach by studying known benign and pathogenic variants, then applied the technique to a set of variants that were not previously studied. We observed a range of functional properties among the variants with a substantial fraction exhibiting complex or mixed patterns of dysfunction. Our findings suggest a need to re-evaluate the simple binary classification of *SCN2A* variants and to engage in efforts to determine how variants with mixed properties promote neuronal dysfunction.

## MATERIALS AND METHODS

### Cell Culture

HEK293T cells (CRL-3216, American Type Culture Collection, Manassas, VA, USA) were stably transfected with the human sodium channel β_1_ (SCN1B) and β_2_ (SCN2B) auxiliary subunits facilitated by *piggyBac* transposon mediated genome insertion (13) The resulting cell line (HEK-beta cells) were maintained in Dulbecco’s modified Eagle’s medium (GIBCO/Invitrogen, San Diego, CA, USA) supplemented with 10% fetal bovine serum (Atlanta Biologicals, Norcross, GA, USA), 2 mM L-glutamine, 50 units/mL penicillin, and 50 µg/mL streptomycin at 37°C in 5% CO_2_.

### Plasmids and mutagenesis

Full-length cDNAs encoding WT or variant intron-stabilized human Na_V_1.2 corresponding to both adult (NCBI accession NM_021007) and neonatal (NCBI accession NM_001371246) splice isoforms (14, 15) were engineered into vectors in which we introduced a high efficiency encephalomyocarditis virus internal ribosome entry site (IRES) with A6 bifurcation sequence (16) followed by the monomeric red fluorescent protein mScarlet (pIRES2-mScarlet).

Variants were introduced into Na_V_1.2 by site-directed mutagenesis using Q5 2X high-fidelity DNA polymerase Master Mix (New England Biolabs, Ipswich, MA) as previously described (15). Mutagenic primers were designed using custom software (available upon request) and are presented in **Table S1**.

### Electroporation

For automated electrophysiology experiments, plasmids encoding WT or variant Na_V_1.2 were electroporated into HEK-beta cells using the MaxCyte STX system (MaxCyte Inc., Gaithersburg, MD, USA) (17). Cells were grown to 70-80% confluence and harvested using TrypLE (ThermoFisher, Waltham, MA). A 500 µL aliquot of cell suspension was used to determine cell number and viability using an automated cell counter (ViCell, Beckman Coulter, Brea, CA, USA). Remaining cells were collected by gentle centrifugation (193 x g, 4 min) at room temperature, followed by washing the cell pellet with 5 mL electroporation buffer (EBR100, MaxCyte Inc.). Cells were resuspended at a final density of 10_8_ viable cells/mL.

For electroporations, 50 µg of WT or variant Na_V_1.2 was mixed with 100 µL of cell suspension (10_8_ cells/mL). The DNA-cell suspension mix was then transferred to an OC-100×2 processing assembly and electroporated using the Optimization 4 protocol. Immediately after electroporation, 10 µL DNase I (Sigma-Aldrich) was added to the DNA-cell suspension mix, and the entire mixture was transferred to a 60 mm dish and incubated for 30 minutes at 37°C in 5% CO_2_. Following incubation, cells were gently resuspended and grown in a T75 flask for 48 hours at 37°C in 5% CO_2_. Cells were then harvested, transfection efficiency determined by flow cytometry (see below), and then frozen in 1 mL aliquots at 1.8 × 10_6_ viable cells/mL.

For manual patch clamp recording experiments, HEK-beta cells were transiently transfected with WT or variant Na_V_1.2 (2 µg) using Qiagen SuperFect reagent (Qiagen, Valencia, CA, U.S.A.).

### Flow Cytometry

Transfection efficiency following electroporation was assessed prior to cell freezing using a BD FACSCanto II flow cytometer (BD Biosciences, Franklin Lake, New Jersey, USA). Forward scatter (FSC), side scatter (SSC), and red fluorescence (PE) were recorded. A 561 nm excitation laser was used. FSC and SSC were used to gate single viable cells and eliminate doublets, dead cells, and debris. Ten thousand events were recorded for each sample, and non-transfected HEK-beta cells were used as a negative control. Data were analyzed using BD FACSDiva 8.0.2.

### Cell preparation for automated electrophysiology

Electroporated cells were thawed one day before experiments, and grown ∼24 hours at 37_°_C in 5% CO2. Prior to experiments, cells were dispersed using TrypLE, and a 500 µL aliquot was taken to determine cell number and viability by automated cell counting. Cells were gently centrifuged (193 × g) for 4 min at room temperature and resuspended at a final density of 180,000 viable cells/mL in external recording solution (see below). Cells were allowed to recover on a shaking rotating platform (200 rpm) at 15°C for 15-30 minutes prior to recording.

### Automated electrophysiology

Automated patch clamp recording was performed using the Nanion SyncroPatch 768PE platform (Nanion Technologies, Munich, Germany) (17). Single-hole low resistance (2-3.5 MΩ) recording chips were used for this study. Pulse generation and data collection were performed using PatchControl384 v1.6.6 and DataControl384 v1.6.0 software (Nanion Technologies). Whole-cell currents were acquired at 10 kHz, series resistance was compensated 80%, and leak currents were subtracted using P/4 subtraction. Whole-cell currents were recorded at room temperature using voltage protocols illustrated in **Fig. S1**. The external solution contained (in mM): 140 NaCl, 4 KCl, 2 CaCl_2_, 1 MgCl_2_, 10 HEPES, 5 glucose, with the final pH adjusted to 7.4 with NaOH, and osmolality adjusted to 300 mOsm/kg with sucrose. The composition of the internal solution was (in mM): 110 CsF, 10 CsCl, 10 NaCl, 20 EGTA, 10 HEPES, with the final pH adjusted to 7.2 with CsOH, and osmolality adjusted to 300 mOsm/kg with sucrose. High resistance seals were obtained by addition of 10 µl seal enhancer solution comprised of (in mM): 125 NaCl, 3.75 KCl, 10.25 CaCl_2_, 3.25 MgCl_2_, 10 HEPES, final pH adjusted to 7.4 with NaOH, followed immediately by addition of 30 µl of external solution to each well. Prior to recording, cells were washed twice with external solution, and the final concentrations of CaCl_2_ and MgCl_2_ were 3 mM and 1.3 mM, respectively. Stringent criteria were used to select cells for inclusion in the final analysis (seal resistance ≥ 200 MΩ, access resistance ≤ 20 MΩ, capacitance ≥ 2 pF, and sodium reversal potential between 45 and 85 mV. All biophysical data were collected from cells whose currents were larger than -200 pA. Voltage control was assessed from conductance-voltage (GV) curves and cells were included in the final analysis if two adjacent points on the GV curve showed no more than a 7-fold increase. Unless otherwise noted, all chemicals were obtained from SigmaAldrich (St. Louis, MO, USA).

A typical experiment recorded from cells expressing WT-Na_V_1.2 and either five Na_V_1.2 variants or four Na_V_1.2 variants plus non-transfected cells seeded into 64-well clusters of a 384-well patch clamp plate. Cells recorded at the same time were electroporated with either WT or variant plasmids in parallel on the same day. To ensure that sufficient numbers of cells were recorded to account for attrition from stringent quality control data filters, we recorded from two 384-well plates simultaneously. Because both plates were run simultaneously, and the plate layout was identical, we combined data from both plates and normalized the data for each variant to the average WT values recorded on the same day. Biophysical properties were listed as not-determined (ND) if less than 5 replicates were obtained for that property for any given variant.

### Manual patch clamp recording

Whole-cell voltage-clamp experiments of heterologous cells were performed as previously described (14). Recordings were made at room temperature using an Axopatch 200B amplifier (Molecular Devices, LLC, Sunnyvale, CA, USA). Patch pipettes were pulled from borosilicate glass capillaries (Harvard Apparatus Ltd., Edenbridge, Kent, UK) with a multistage P-1000 Flaming-Brown micropipette puller (Sutter Instruments Co., San Rafael, CA, USA) and fire-polished using a microforge (Narashige MF-830; Tokyo, JP) to a resistance of 1.5–2.5 MΩ. The pipette solution consisted of (in mM): 110 CsF, 10 CsCl, 10 NaCl, 20 EGTA, 10 HEPES, with the final pH adjusted to 7.2 with CsOH, and osmolality adjusted to 300 mOsm/kg with sucrose. Cells in the recording chamber were superperfused with bath solution containing (in mM): 140 NaCl, 4 KCl, 3 CaCl_2_, 2 MgCl_2_, 1 HEPES, 5 glucose, with the final pH adjusted to 7.4 with NaOH, and osmolality adjusted to 300 mOsm/kg with sucrose.

### Data Analysis

Data were analyzed and plotted using a combination of DataControl384 v1.6.0 (Nanion Technologies), Clampfit 10.4 (Molecular Devices), Microsoft Excel (Microsoft Office 2019), and GraphPad Prism (GraphPad Software, San Deigo, CA, USA). Whole-cell currents were normalized to membrane capacitance, and data are expressed as mean ± SEM unless otherwise noted. GraphPad Prism was used to fit voltage-dependence of activation and inactivation curves with Boltzmann functions, and recovery from inactivation with a 2-exponential equation. Window current was calculated by integrating the area under the intersection between the Boltzmann fits for voltage-dependence of activation and inactivation using a custom MatLab script (18). All data were normalized to WT run in parallel and data were treated as non-parametric for statistical analyses. Statistical comparison of manual and automated recording of Nav1.2A-WT was performed using a Mann-Whitney test. Statistical analyses of Nav1.2 variants were performed using a Kruskal-Wallis test followed by Dunn’s post-hoc test for multiple comparisons. The threshold for statistical significance was p ≤ 0.05.

## RESULTS

### Automated electrophysiological analysis of Na_V_1.2

We employed a high-efficiency electroporation method to transiently express recombinant human Na_V_1.2 suitable for automated electrophysiology. The Na_V_1.2 subunit was coupled to mScarlet expression, and electroporation efficiency was quantified by flow cytometry. We constructed separate expression plasmids encoding either the canonical adult brain-expressed splice variant (Na_V_1.2A) or one with an alternative exon 5 predominantly expressed in developing brain (neonatal Na_V_1.2; Na_V_1.2N). Transfection efficiencies of 60-80% were achieved in HEK293T cells stably transfected with β_1_ and β_2_ accessory subunits.

We recorded robust whole-cell sodium currents using a dual 384-well automated electrophysiology platform and compared results with experiments performed on the same cells using manual patch clamp recording (**Fig. S2A, B**). The average peak current density measured by automated patch-clamp was smaller than that obtained by manual patch clamp (**Fig. S2B**), but normalized current-voltage relationships were nearly identical between the two recording methods (**Fig. S2C**). The voltage-dependence of activation was not significantly different between recording methods (**Fig. S2D**; **Table S2**), whereas other parameters showed small differences between recording methods (**Fig. S2D, E, F; Table S2**). To control for non-specific factors such as cell passage number, we designed automated patch clamp experiments to compare Na_V_1.2 variants only to the WT channel electroporated and assayed simultaneously. Electrophysiological data from 5,858 cells were analyzed for this study.

### Analytical validity of automated patch clamp for Na_V_1.2

To demonstrate analytical validity of automated patch clamp for Na_V_1.2, we engineered a collection of nonsynonymous population variants (R19K, K908R, E1153K, and G1522A) and previously published pathogenic missense variants (R853Q, R937C, M1879T, and R1882Q) in Na_V_1.2A to serve as a validation set (**Fig. 1; Fig. S3**). Population variants were chosen based on minor allele frequency greater than 0.0001 in gnomAD (19), while pathogenic variants were chosen to represent either gain- and loss-of-function effects based on previously published data (10, 18, 20-22).

**Fig. 1.**
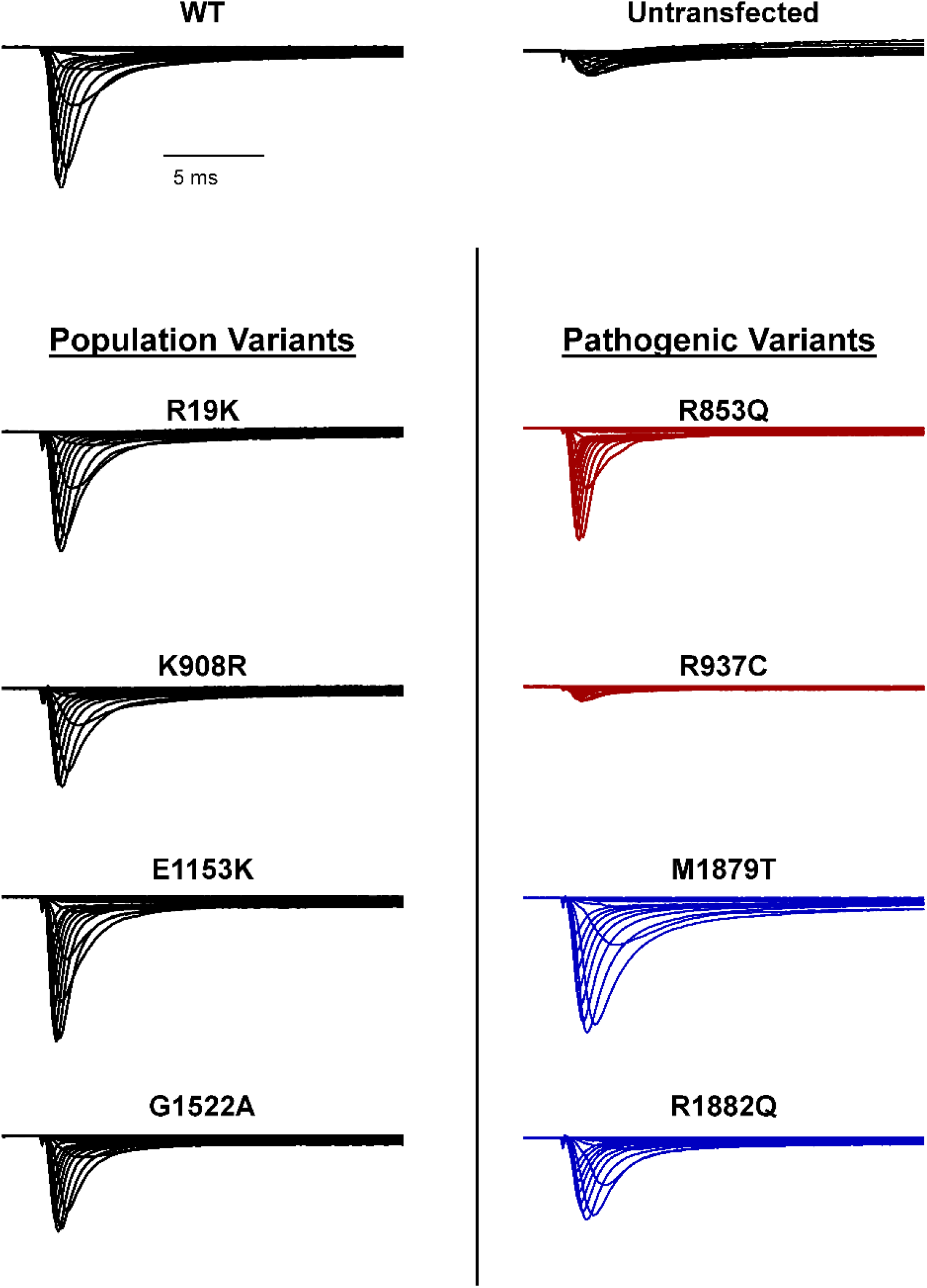
Functional validation of a training set of Na_V_1.2 variants. Average normalized whole-cell sodium currents of (A) WT Na_V_1.2 (left), untransfected cells (right), **(B)** a set of population variants, and **(C)** known pathogenic variants representing loss-of-function (LOF, red) -and gain-of-function (GOF, blue) phenotypes. Average traces are from 5 to 65 cells.

We measured several biophysical properties (**Fig. S1**) to fully capture the spectrum of sodium channel function, including whole-cell current density, voltage-dependence of activation and inactivation, recovery from inactivation, frequency-dependent channel rundown, time-constant for onset of inactivation at 0 mV, charge movement in response to a voltage ramp, and persistent sodium current. For the population variants, only G1522A showed a significant difference in whole-cell sodium current density compared to WT channels (fraction of WT current: 0.69 ± 0.9, n=25, p=0.0278). For all other parameters R19K, K908R, E1153K, and G1522A were indistinguishable from WT-Na_V_1.2 (**Fig. 1, Table S3**).

Among the pathogenic variants chosen for the validation set we chose two previously classified loss-of-function variants (R853Q, R937C) and two gain-of-function variants (M1879T, R1882Q). Our analysis of these variants using automated electrophysiology largely recapitulated previously published results with the exception of R853Q current density, which was not significantly different from WT channels (**Table S3**), whereas other properties conferring loss-of-function were consistent with previous reports (18, 20). For the other loss-of-function variant, R937C, whole-cell sodium current (0.18 ± 0.02 of WT; **Fig. 1, Table S3**) was indistinguishable from current recorded from non-transfected cells (0.17 ± 0.01 of WT, **Table S3**). Automated patch clamp recordings of the gain-of-function variants M1879T and R1882Q in the validation set demonstrated similar patterns of dysfunction including depolarized shifts in the voltage-dependence of inactivation, slower inactivation time course, and a larger ramp current as reported previously (**Fig. 1, Table S3**). Results from the validation set demonstrate that we can use automated patch clamp to measure multiple gain- and loss-of-function properties for pathogenic *SCN2A* variants and that non-pathogenic population variants are benchmarks for normal Na_V_1.2 function.

### Functional analysis of disease-associated Na_V_1.2 variants

We used this optimized approach to determine the functional consequences of 28 Na_V_1.2 variants identified through a patient registry established at The Ann and Robert H. Lurie Children’s Hospital of Chicago (**Fig. S3**) (*manuscript in preparation*). This list included three of the previously mentioned pathogenic variants (R853Q, M1879T, R1882Q) that are part of the validation set. Most of the variants (n = 22) were associated with developmental and epileptic encephalopathy (DEE) with seizure onset within the first month of life, whereas five were associated with DEE having seizure onset after 6 months of age while one variant was associated with impaired neurodevelopment without seizures. One variant (G211D) affected a residue within the neonatal-expressed exon 5N, and therefore was only studied in Na_V_1.2N. By contrast, E1211K, K1422S, S1758R, and A1773T were only studied in the canonical adult splice isoform because of the associated late clinical onset. All other variants were studied in both adult and neonatal Na_V_1.2 splice isoforms.

Ten of the analyzed variants expressed in the canonical Na_V_1.2A isoform exhibited significantly smaller peak current density than WT (**Fig. 2, Figs. S4, Table S4**), ranging from a modest reduction (A880S), to complete loss of function (F978L; **Fig. 2B, Table S4**). Two variants, D1050V and R1626Q, had current densities larger than WT (**Fig. 2B, Table S4**). We observed significant differences in the voltage-dependence of activation for seven variants with four exhibiting a hyperpolarized shift (A427D, G879R, E1211K, and K1260Q), and three a depolarized shift (R571H, R1882L, and R1882Q; **Fig. 3, Table S4**). Voltage-dependence of inactivation was significantly different from WT channels for 20 variants including 12 variants with hyperpolarized shifts (**Fig. 4, Table S4**). Interestingly, there is a cluster of variants in domain IV and the C-terminus that all exhibited depolarized shifts in the voltage-dependence of inactivation (I1537S/M1538I, R1626Q, S1780I, M1879T, R1882L, and R1882Q; **Fig. 4, Table S4**).

**Fig. 2.**
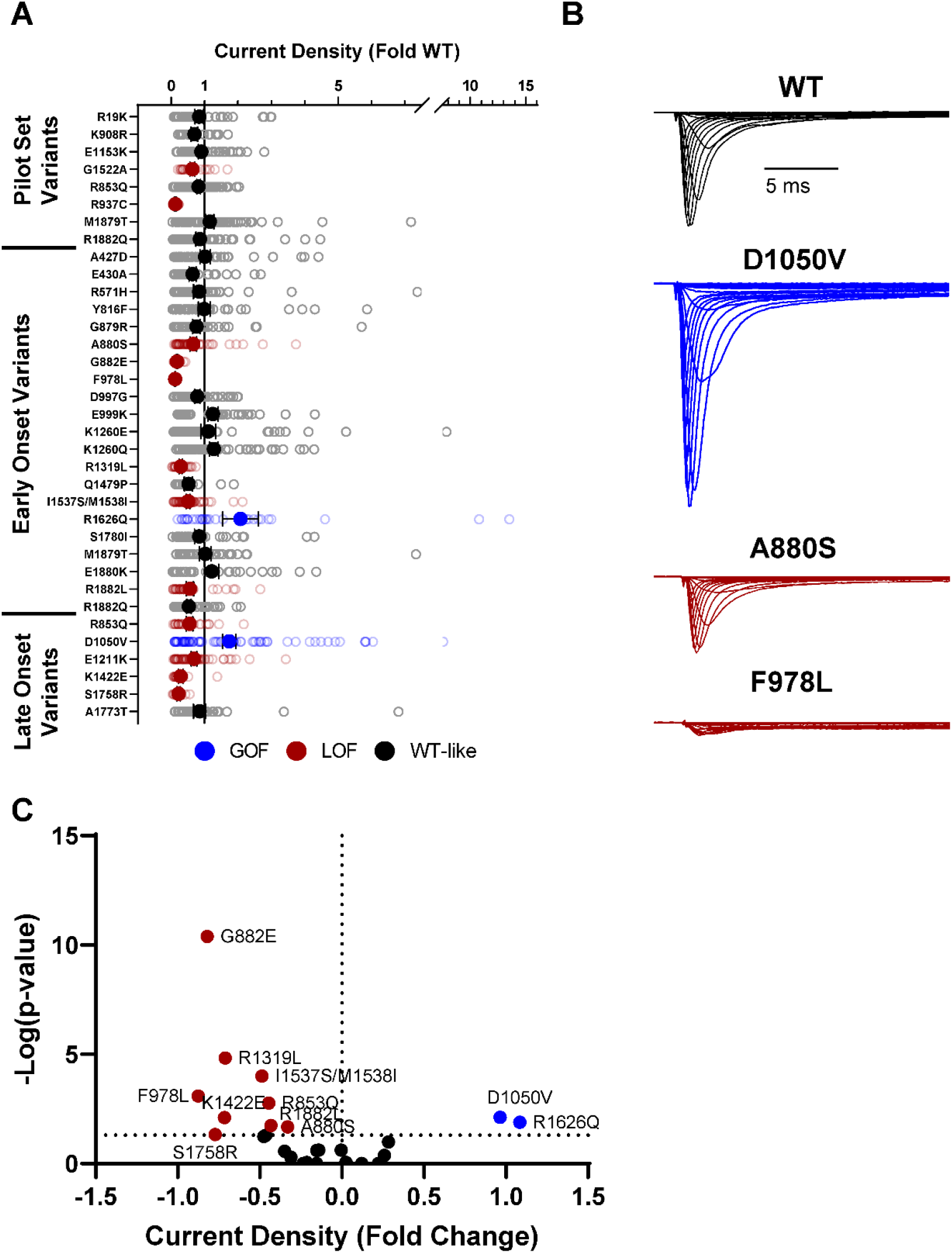
Na_V_1.2 variants alter whole-cell current density. **(A)** Average deviation of whole-cell sodium current density from WT Na_V_1.2 for population and disease-associated variants. **(B)** Average traces for WT Na_V_1.2, a gain-of-function (GOF) variant Na_V_1.2A-D1050V, and two loss-of-function (LOF) variants Na_V_1.2A-A880S, and Na_V_1.2A-F978L. **(C)** Volcano plot highlighting variants significantly different from WT. Red symbols denote loss-of-function and blue symbols denote gain-of-function with p < 0.05 (n = 5-79).

**Fig. 3.**
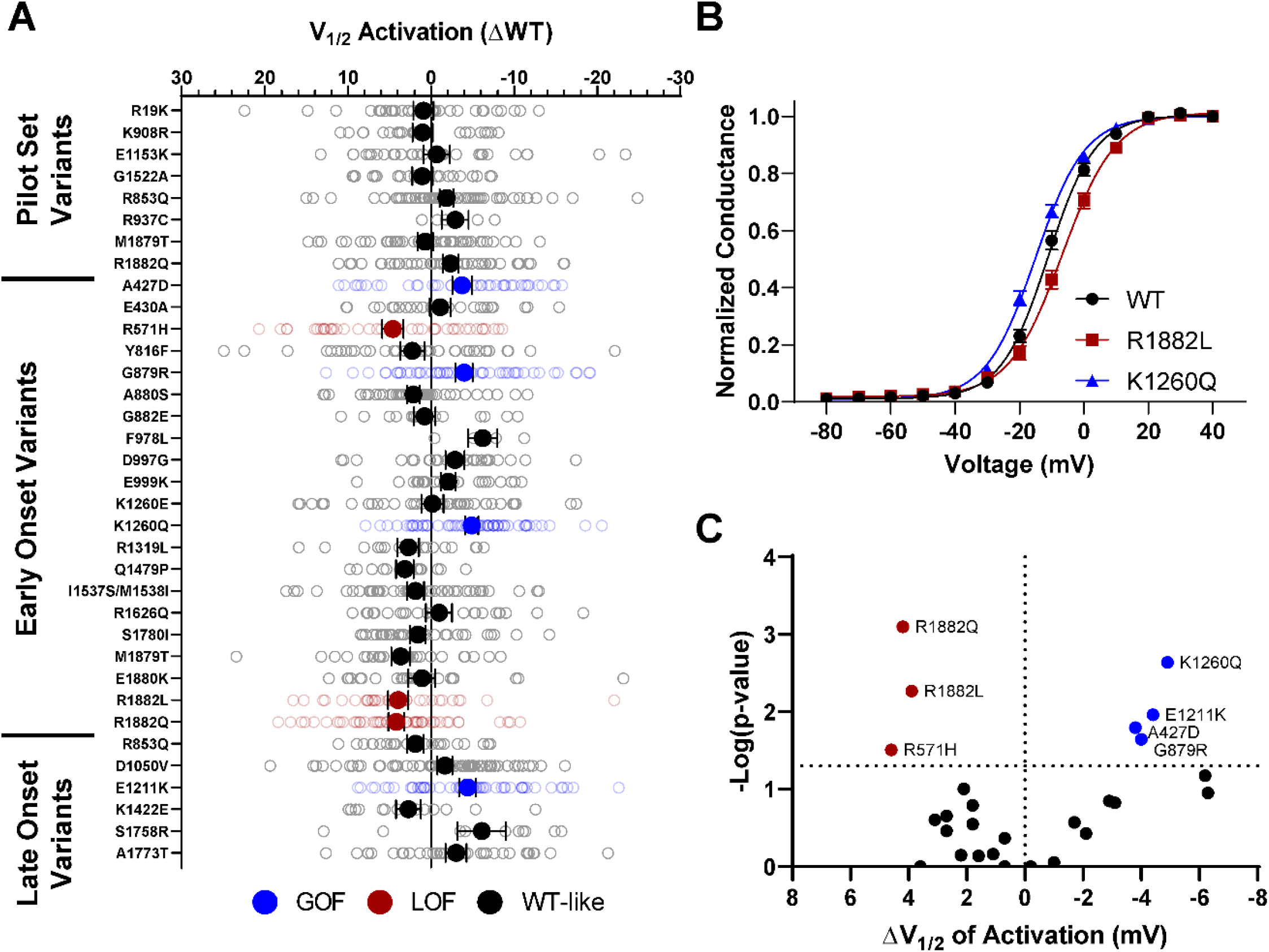
Na_V_1.2 variants alter voltage-dependence of activation. **(A)** Average deviation from WT Na_V_1.2 for V_1/2_ of activation (in mV). **(B)** Conductance-voltage curves showing a gain-of-function (Na_V_1.2-K1260Q) and a loss-of-function (R1882L) variant. **(C)** Volcano plot highlighting variants significantly different from WT. Red symbols denote loss-of-function and blue symbols denote gain-of-function with p < 0.05 (n = 5-65).

**Fig. 4.**
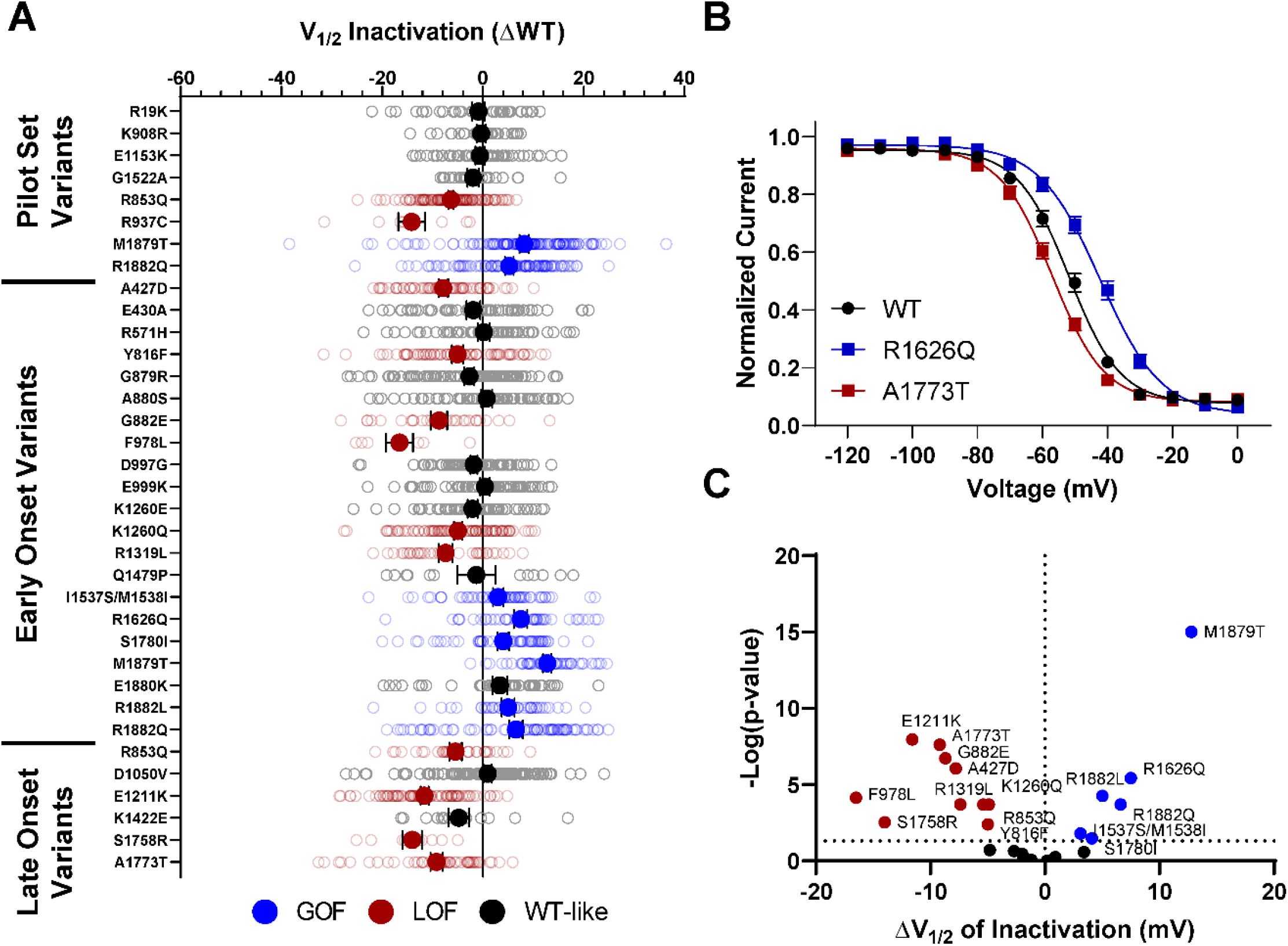
Na_V_1.2 variants alter voltage-dependence of inactivation. **(A)** Average deviation from WT Na_V_1.2 for V_1/2_ of inactivation (in mV). **(B)** Steady-state inactivation curves showing a gain-of-function (Na_V_1.2-R1626Q) and a loss-of-function (Na_V_1.2-A1773T) variant. **(C)** Volcano plot highlighting variants significantly different from WT. Red symbols denote loss-of-function and blue symbols denote gain-of-function with p < 0.05 (n = 8-122).

Voltage-dependence of activation and inactivation curves overlap to define a region called the window current where channels are open and not inactivated. The presence of larger or smaller window currents may contribute to loss- or gain-of-function driven by differences in voltage-dependent gating. Variants with larger depolarization of voltage-dependence of inactivation such as I1537S/M1538I, R1626Q, M1879T, R1882L, and R1882Q, exhibited larger window current compared to WT channels (**Fig. S5A,C, Table S4**). Similarly, R571H, which has a hyperpolarized voltage-dependence of activation, also shows larger window current compared to WT. Interestingly, E1211K shows both a hyperpolarization of activation and inactivation. The resulting window current is smaller compared to WT and suggests that E1211K may result in an overall loss-of-function with respect to voltage-dependent gating (**Fig S5A D, Table S4**).

Domain IV in Na_V_ channels contributes to the voltage-dependence of channel activity and to the time course for the onset of channel inactivation. Thus, in addition to shifts in voltage-dependence of inactivation, variants in domain IV and the C-terminus may slow the onset of channel inactivation, which can lead to a slower time course of inactivation (reflected by larger time constants) and larger currents during a depolarizing voltage ramp protocol (**Fig. S1**). Indeed, seven of the 11 variants with slower inactivation kinetics reside in these domains (I1537S/M1538I, R1626Q, A1773T, M1879T, E1880K, R1882L, and R1882Q; **Fig. 5, Table S4**). Four of these variants (R1626Q, E1880K, M1879T, and R1882L) also showed larger ramp currents compared to WT (**Fig. 6, Table S4**). One variant, G879R, exhibited a larger ramp current compared to WT, even though the onset of inactivation was slightly faster than WT. However, this larger ramp current can be attributed to a larger persistent sodium current compared to WT (**Fig. S6**). We also measured frequency dependent rundown (**Fig. S6**), and recovery from inactivation (**Fig. S7**) for each variant. While some variants showed gain-of-function effects, loss-of-function was the predominant phenotype for recovery from inactivation and frequency dependent rundown.

**Fig. 5.**
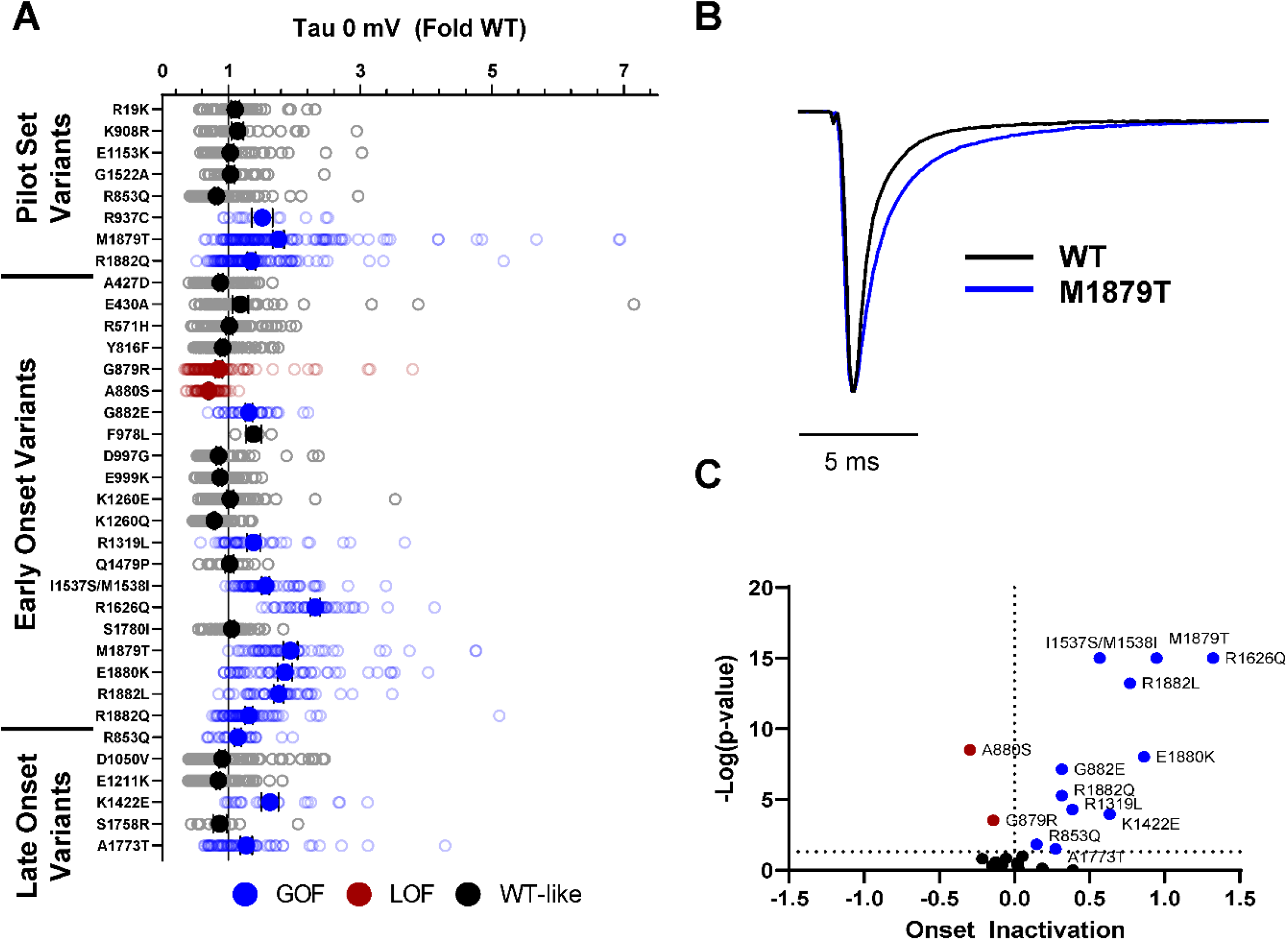
Na_V_1.2 variants alter inactivation time constants. **(A)** Average deviation of inactivation time constant (tau) measured at 0 mV from WT Na_V_1.2 for disease-associated variants. **(B)** Average traces for WT Nav1.2 and a GOF variant Na_V_1.2-M1879T recorded at 0 mV. **(C)** Volcano plot highlighting variants significantly different from WT. Red symbols denote loss-of-function and blue symbols denote gain-of-function with p < 0.05 (n = 9-120).

**Fig. 6.**
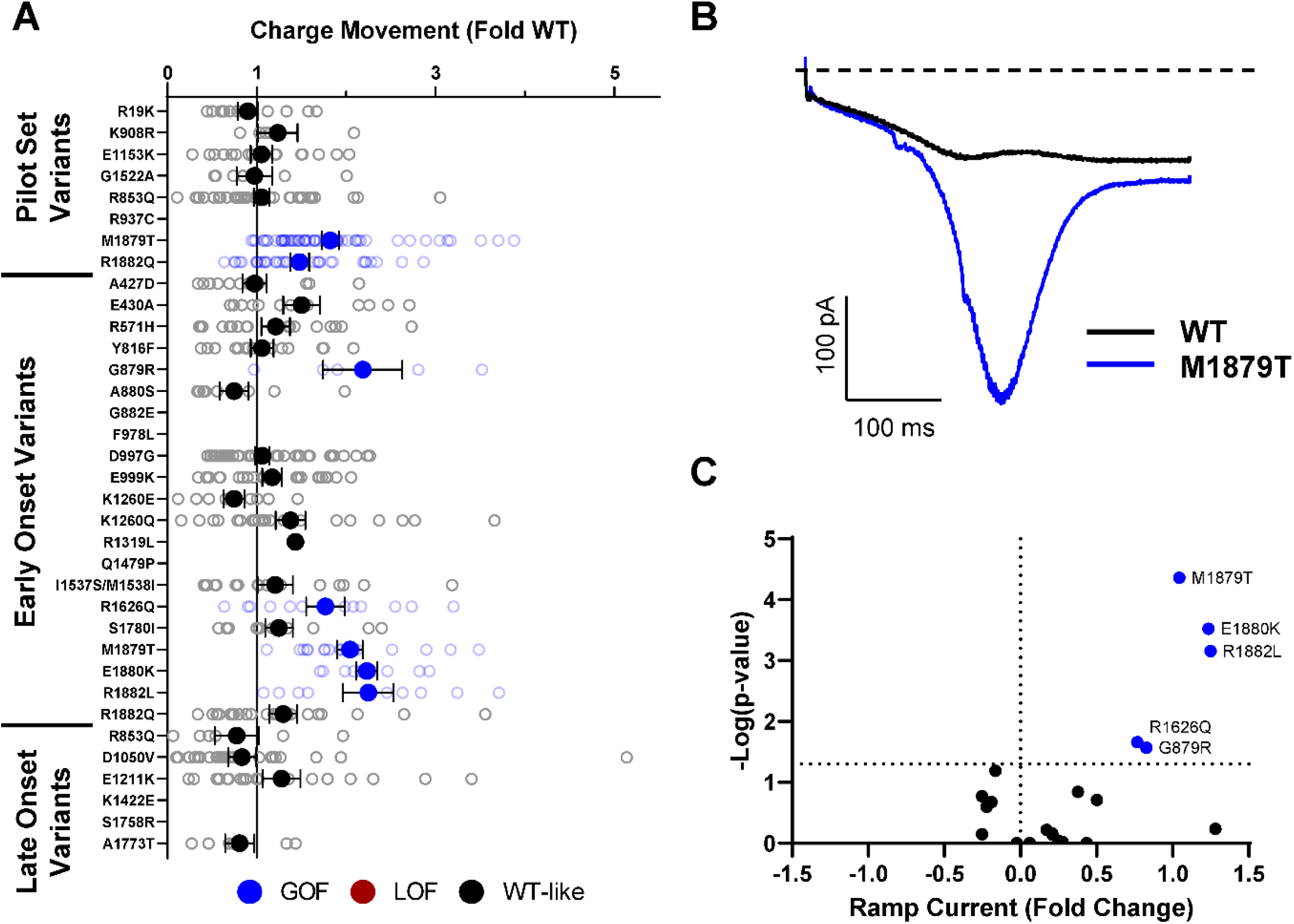
Na_V_1.2 variants alter inactivation ramp currents. **(A)** Average deviation of ramp currents from WT Na_V_1.2 for disease-associated variants. **(B)** Average traces for WT Na_V_1.2 and a GOF variant Na_V_1.2-M1879T. **(C)** Volcano plot highlighting variants significantly different from WT. Red symbols denote loss-of-function and blue symbols denote gain-of-function with p < 0.05 (n = 5-45).

A small number of variants were challenging to evaluate using automated patch clamp due to either poor viability after electroporation, or very small whole-cell currents that could not be distinguished from endogenous currents. We used manual patch clamp recording to validate the phenotypes of three variants, G882E, F978L, and Q1479P. Both G882E and F978L exhibited very small currents compared to WT when recorded using either automated or manual patch clamp (**Fig. 7A B**). Although whole-cell current amplitude was small for G882E, they were large enough to reliably measure voltage-dependence of activation and inactivation, and similar results were obtained from automated and manual patch clamp recording. Voltage-dependence of activation for G882E was similar to that of WT (WT: -21.5 ± 1.2 mV, n=19, G882E: -21.1 ± 1.9 mV, n = 10, p=0.9073), while exhibiting a hyperpolarized shift in voltage-dependence of inactivation (WT: - 58.4 ± 0.9 mV, n=19, G882E: -63.3 ± 2.0 mV, n = 10, p=0.0193, **Fig. 7B**).

**Fig. 7.**
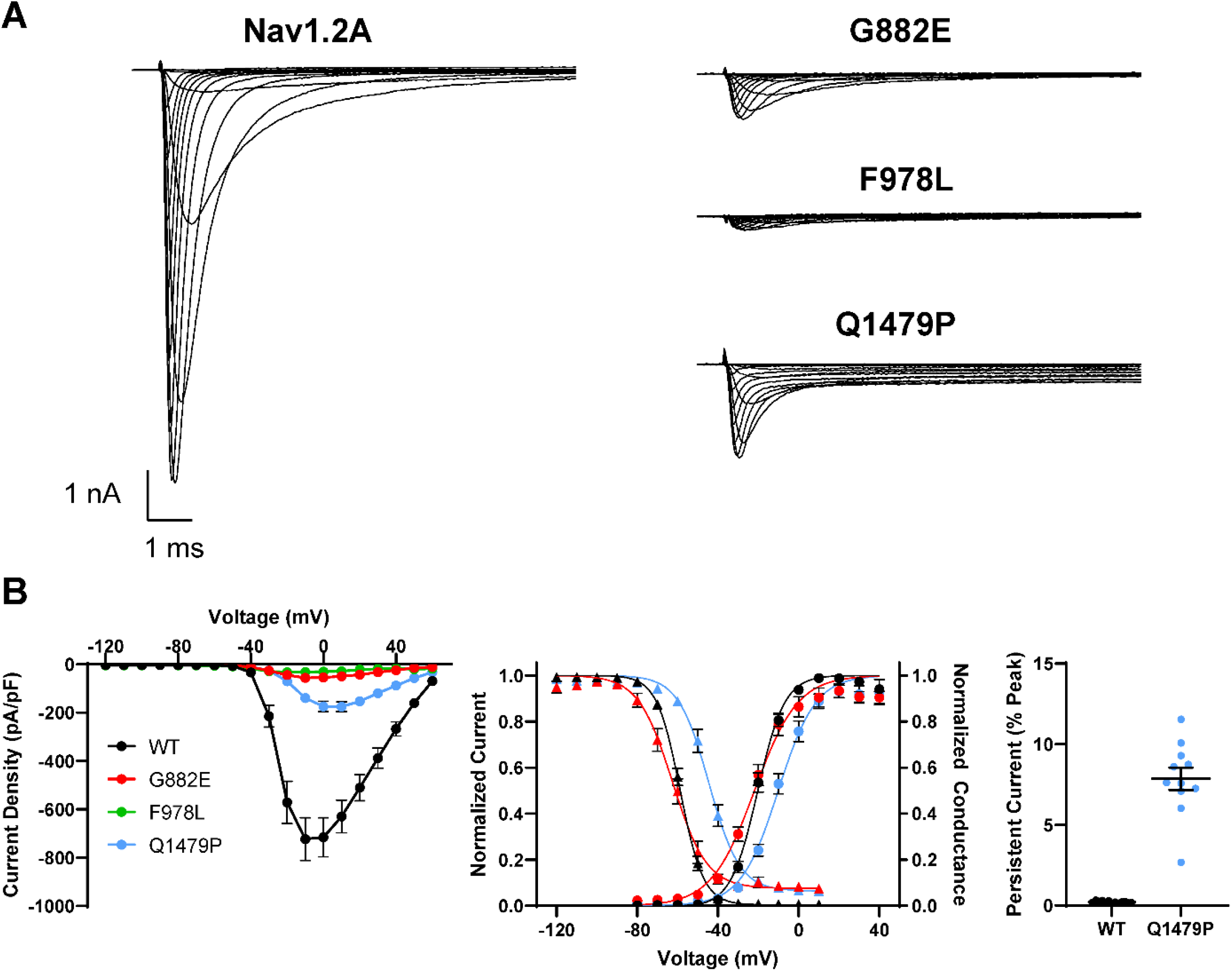
Validation of difficult to record Na_V_1.2 variants using manual patch clamp. **(A)** Average whole-cell current traces of WT and three difficult to characterize variants (G882E, F978L, and Q1479P) recorded using manual patch-clamp. **(B)** Summary current voltage relationship (*left*), voltage-dependence of activation and inactivation (*middle*), and persistent current (*right*) of Na_V_1.2 variants recorded using manual patch clamp. All data were from 9 to 19 cells.

Cells electroporated with Q1479P had poor viability compared to cells electroplated with WT Na_V_1.2. Automated patch clamp recordings showed smaller currents than WT, but indicated a large persistent sodium current that could potentially be toxic to cells. Manual patch clamp recording revealed that while the peak whole-cell sodium currents were indeed smaller than that of WT, persistent current was 38 times larger than WT when measured as a percentage of peak current (WT: 0.2 ± 0.02%, n=12, Q1479P: 7.7 ± 0.8%, n=9, p<0.00001, **Fig. 7B**) and 8-fold larger than WT when measured as current density (WT: -2.3 ± 0.7 pA/pF, Q1479P: -18.2 ± 4.9 pA/pF, p=0.0048). Manual recording of Q1479P also revealed depolarized shifts in voltage-dependence of activation (WT: -21.5 ± 1.2 mV, n=19; Q1479P: -10.2 ± 1.7 mV, n=11, p<0.0001) and voltage-dependence of inactivation (WT: -58.4 ± 0.9 mV, n=19; Q1479P: -44.2 ± 1.6 mV, n=11, p<0.0001, **Fig. 7B**)

### Comparison of adult and neonatal splice isoforms

We studied variants associated with early-onset DEE in the neonatal Na_V_1.2N splice isoform (**Figs. S8-S14, Table S5**), and found a mix of gain-and loss-of-function properties, similar to that observed for the variants expressed in the canonical Na_V_1.2A isoform. There are a few variants for which splice isoform impacts the functional properties differently. For example, both R571H and K1260E expressed in the neonatal isoform exhibited hyperpolarized voltage-dependence of activation compared to WT (**Fig. S14**), whereas expression in Na_V_1.2A was associated with a depolarized or WT-like activation voltage-dependence for R571H and K1260E, respectively. Additionally, while R853Q exhibited a greater sensitivity to frequency dependent rundown in the adult isoform its behavior was WT-like in the neonatal isoform (**Fig. S14**).

### Integrated summary of Na_V_1.2 variant dysfunction

Assigning the net effects of a variant as either gain- or loss-of-function is challenging when considering the constellation of all measured functional properties. Complete loss-of-function is easy to assign when there is no measurable sodium current (e.g., R937C). Similarly, variants with certain properties such as isolated enhanced persistent current, slower inactivation time course, or a large depolarizing shift in the voltage-dependence of inactivation can be inferred to be gain-of-function. However, most variants we studied exhibited mixed properties that are difficult to categorize according to this simple binary scheme. To provide a visual and integrated summary of functional properties for each variant ascertained in the two different splice variants, we constructed heat maps that scaled each biophysical parameter along a functional axis from loss to gain (**Fig. 8**). When the data are displayed this way, many Na_V_1.2 variants appear to exhibit complex patterns of dysfunction with two or more properties exhibiting opposing effects, which is inconsistent with a simple binary overall functional effect.

**Fig. 8.**
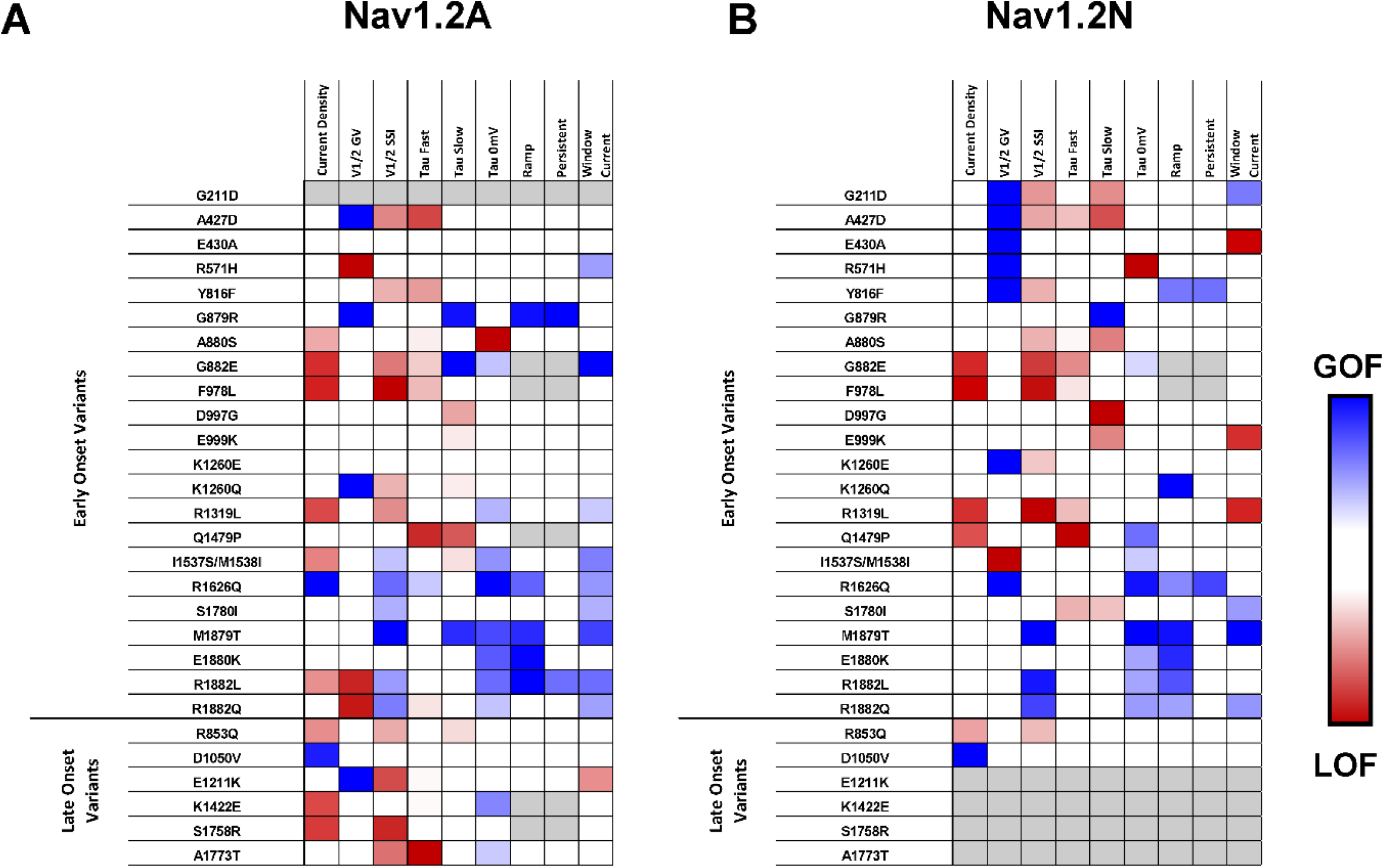
Comparison of epilepsy associated variants in the adult and neonatal isoforms of Nav1.2. Heat maps showing GOF (blue) and LOF (red) phenotypes measured for epilepsy associated Na_V_1.2 variants in the **(A)** adult and **(B)** neonatal splice isoforms. Only properties which reached the threshold for statistical significance are highlighted.

## DISCUSSION

The widespread use of genetic testing in medical practice and research has led to rapid growth in the number of genetic variants identified in ion channel genes associated with monogenic epilepsies. Determining the functional consequences of disease-associated ion channel variants has value in revealing pathogenic mechanisms, contributing to understanding genotype-phenotype relationships, and helping with the assessment of pathogenicity. However, the explosion in genetic data makes the functional evaluation of individual variants using traditional whole-cell patch clamp techniques insufficient, and techniques with more scalable throughput are required to meet demand.

In this study, we demonstrate successful use of automated patch clamp recording to determine the functional properties of several *SCN2A* variants expressed in two splice isoforms. While traditional voltage clamp techniques are considered to be the gold standard for evaluating ion channel function, automated patch clamp offers specific advantages that strengthen experimental rigor including higher throughput, ability to directly compare WT and variants in parallel, and unbiased cell selection (17). Furthermore, the higher throughput achievable with automated patch clamp allows for greater numbers of variants to be studied under standardized experimental conditions thus avoiding inter-laboratory heterogeneity. For this study, we investigated 28 distinct variants with most expressed in two splice isoforms (total 51 constructs) and report data recorded from nearly 6,000 individual cells. This work complements our prior efforts using automated patch clamp to determine the functional consequences and pharmacology of other disease-associated ion channels (23-26).

Our evaluation of a cohort of disease-associated *SCN2A* variants revealed a spectrum of Na_V_1.2 dysfunction that was not easily parsed into the binary categories of gain- and loss-of-function. When considering multiple biophysical parameters, many variants exhibit a constellation of dysfunctional properties that represent mixed gain- and loss-of-function. Good examples of mixed patterns of dysfunction include R1882L and R1882Q. When expressed in Na_V_1.2A, both of variants exhibit depolarizing shifts in activation voltage-dependence combined with slower kinetics of fast inactivation, which represent opposing effects on channel function. Perhaps given the complexity of *SCN2A*-related clinical phenotypes and that physiological function of *SCN2A* is developmentally regulated, complex patterns of variant channel dysfunction is not unexpected. Our finding of mixed functional properties suggests a need for a more nuanced classification of *SCN2A* variant dysfunction. Specifically, our results imply that the gain-versus loss-of-function paradigm oversimplifies variant effect on channel function and does not fully capture the complete physiological impact of channel dysfunction. Our data further emphasize the need to consider which parameters are the key drivers of channel dysfunction and abnormal neuronal physiology. Other scalable experimental approaches such as computational neuronal action potential modeling (27) or dynamic clamp recording (21) may be valuable to determine the net effect of mixed dysfunctional properties on neuronal excitability. While these strategies are beyond the scope of our current study, the data we generated will be valuable to inform these complementary approaches.

Differences in *SCN2A* variant function may segregate with the age of onset of epilepsy or correlate with later onset neurodevelopmental disorders. Among the variants we studied that were associated with early onset seizure disorders, we observed functional properties consistent with an overall gain-of-function (e.g., R1626Q, M1879T), mixed patterns of dysfunction, or overt loss-of-function (e.g., F978L). Similarly, among variants associated with later onset seizures or a neurodevelopmental disorder without seizures (e.g., S1758R) we observed mixed functional properties, although channel loss-of-function was the most prevalent effect (**Fig. 8**). The outlier was D1050V, which exhibited significantly larger peak current density than WT channels without differences in voltage-dependent or kinetic properties. This variant affects a residue in the D2-D3 cytoplasmic domain near an ankyrin binding site. A recent study provided evidence that loss of ankyrin-B scaffolding of Na_V_1.2 in cortical neurons phenocopies the neurophysiological defects observed in *Scn2a* haploinsufficient mice (28). These observations raise the possibility that D1050V disrupts Na_V_1.2 interactions with ankyrin and mimics a channel loss-of-function. Future investigations of the cell surface expression or localization of *SCN2A* variants in neurons are required to address this hypothesis.

*SCN2A* undergoes developmentally regulated alternative mRNA splicing that generated transcripts containing one of two mutually exclusive alternative exons encoding a portion of the first domain voltage-sensor (S3 and S4 helices) with slight differences in amino acid sequence (29, 30). The neonatal expressed Na_V_1.2N splice isoforms predominates during early development whereas transcripts containing the alternative (adult expressed Na_V_1.2A) exon are expressed beginning in the post-natal period (31). We previously reported functional differences between neonatal and adult splice Na_V_1.2 isoforms and pointed out that some epilepsy-associated variants exhibit distinct patterns of dysfunction in the two isoforms (14). Two C-terminal variants (R1882Q, R1882L), which are associated with neonatal onset epileptic encephalopathy, exhibit gain-of-function profiles when expressed in the neonatal isoform but a more mixed profile in the adult isoform.

In summary, we conducted a systematic functional evaluation of a large cohort of disease-associated *SCN2A* variants using an optimized automated patch clamp recording strategy. The scale of this work allowed us to compare the function of variants associated with different disease phenotypes and to examine how functional properties vary between developmentally regulated splice isoforms. Our findings illustrate a prevalence of complex biophysical effects for many variants suggesting that a simple binary scheme to classify variants as either gain- or loss-of-function is insufficient to capture the full spectrum of variant channel dysfunction. The higher throughput capabilities of automated patch clamp enables greater standardization and more robust experimental rigor that is well suited to assessing *SCN2A* variant pathogenicity.

## ACKNOWLEDGEMENTS

This work was supported NIH grant U54 NS108874 (ALG), and a Director’s Award from the Simon’s Foundation Autism Research Initiative (SFARI). This work was supported by the Northwestern University Interdepartmental ImmunoBiology Flow Cytometry Core Facility.

## DISCLOSURES

A.L.G. received research grant funding from Praxis Precision Medicines and Neurocrine Biosciences, and received research grant funding from and is a member of the Scientific Advisory Board for Tevard Biosciences.

## AUTHORS’ CONTRIBUTIONS

C.H.T., F.P., T.V.A., J.M.D., N.F.G. and C.G.V. performed experimental work and analyzed data. J.J.M. identified SCN2A variants. C.H.T. and A.L.G. wrote the manuscript. A.L.G. acquired research funding. All authors reviewed and approved the final version of the manuscript prior to submission.

## SUPPLEMENTAL INFORMATION

### Supplemental Figures

**Fig. S1.**
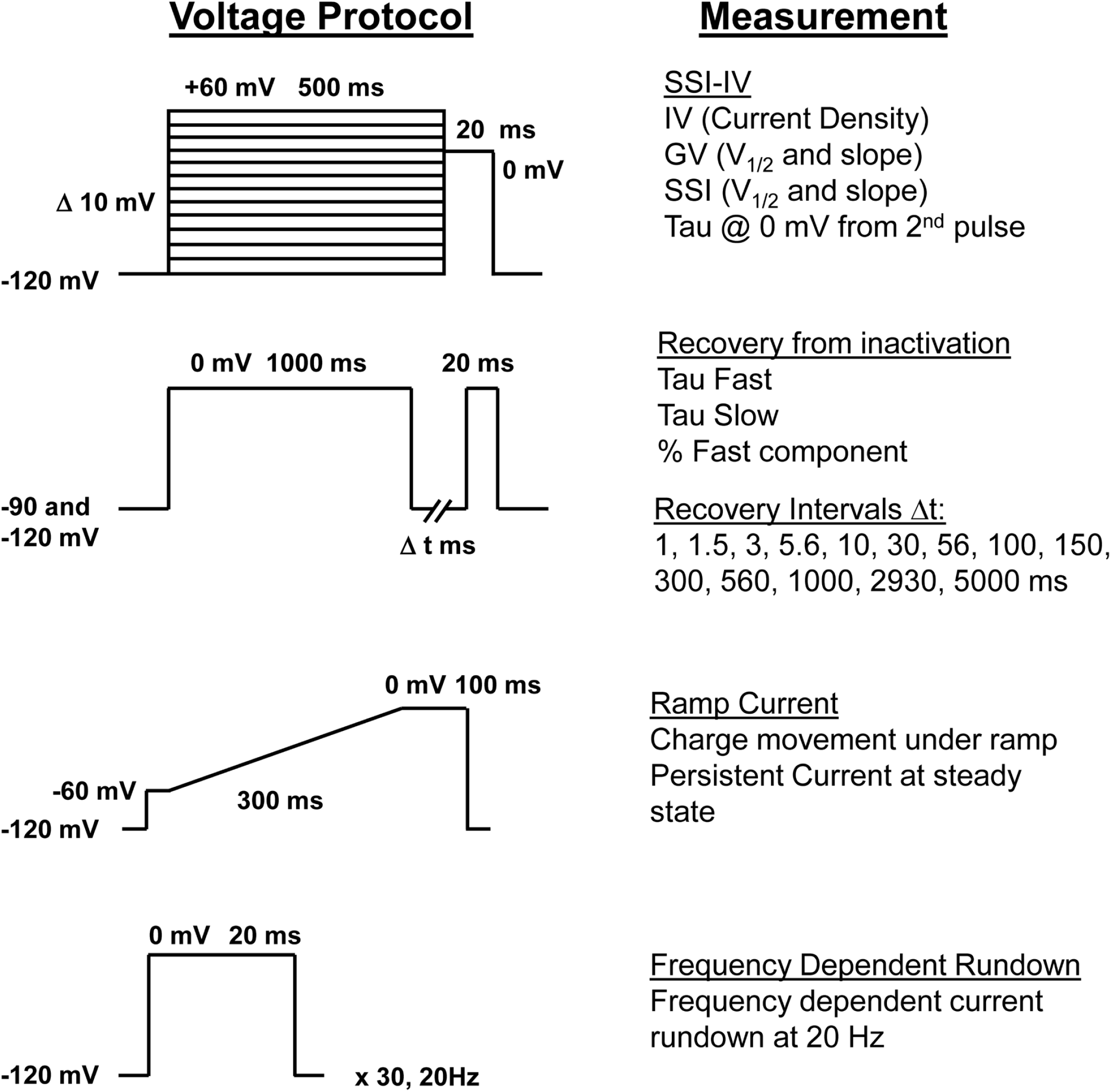
Voltage protocols used to assess voltage gated sodium channel biophysical parameters by both automated and manual electrophysiology.

**Fig. S2.**
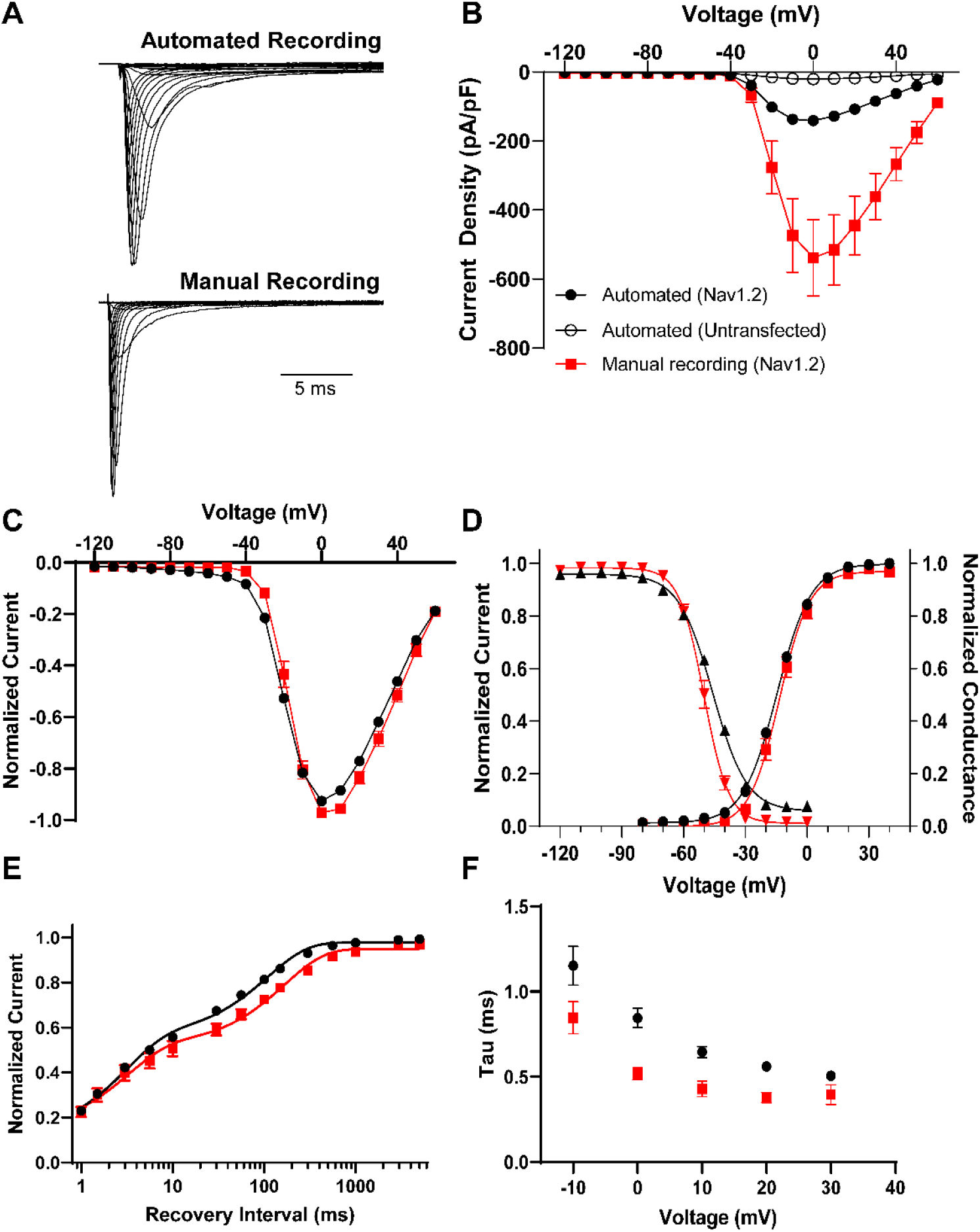
Comparison of automated and manual patch clamp evaluation of Na_V_1.2. **(A)** Average normalized whole-cell sodium currents from automated (top) and manual (bottom) patch clamp recording of cells expressing wild-type (WT) Na_V_1.2. **(B)** Summary current-voltage relationships for automated recording of untransfected cells (open circles), automated recording of cell expressing Na_V_1.2 (black circles), and manual recording of cells expressing Na_V_1.2 (red squares). **(C)** Normalized whole-cell sodium currents comparing automated (black circles) and manual (red squares) recording methods. **(D)** Voltage-dependence of activation and inactivation of Na_V_1.2 comparing automated (black) and manual (red) recording methods. **(E)** Recovery from inactivation of Na_V_1.2 comparing automated (black circles) and manual (red squares) recording methods. **(F)** Time-constant for the onset of inactivation of Na_V_1.2 as a function of membrane potential for automated (black circles) and manual (red squares) recording methods. All data are expressed as mean ± SEM from 14 to 1253 cells.

**Fig. S3.**
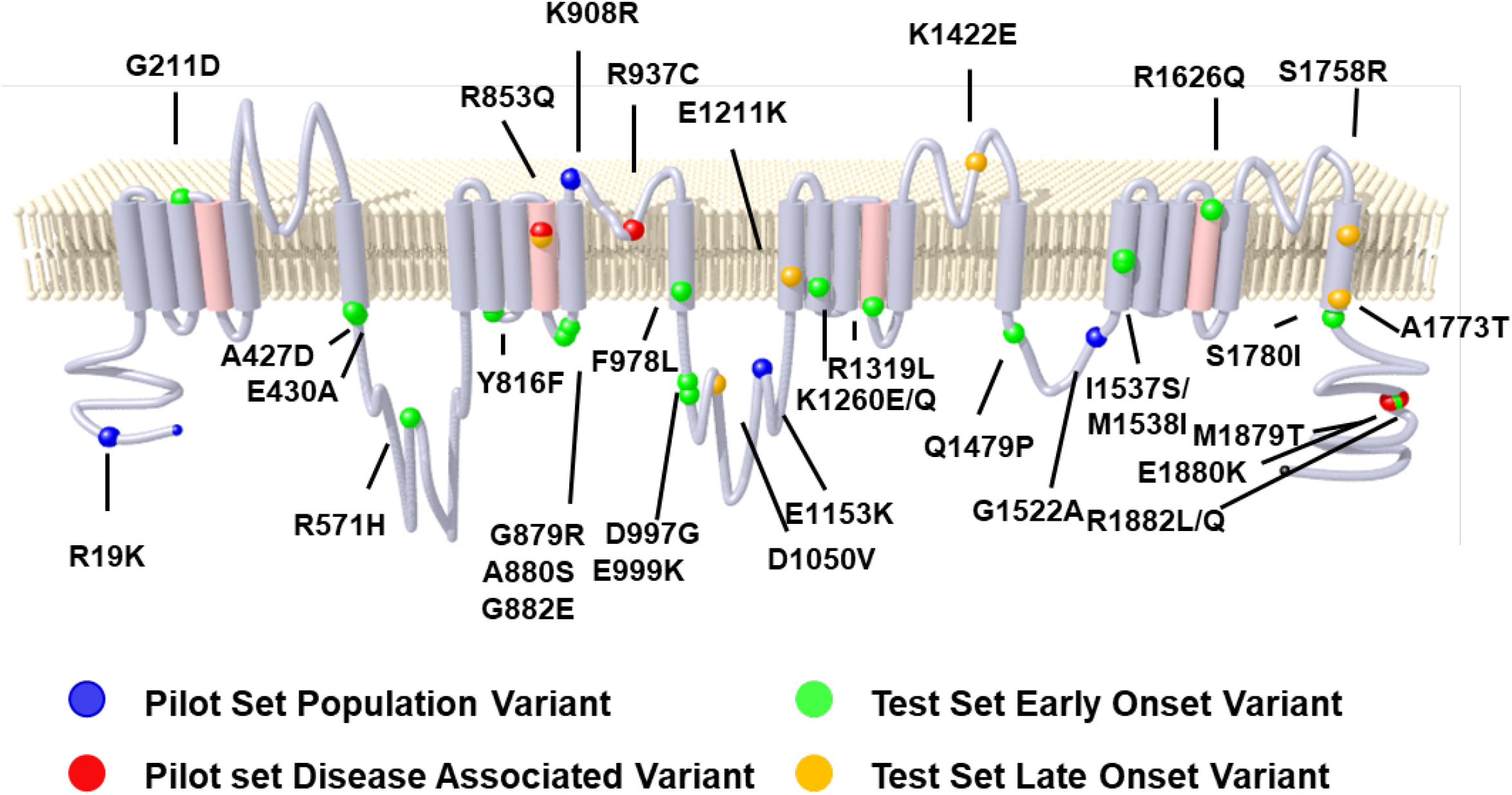
Location of Na_V_1.2 variants. Topology diagram of Na_V_1.2 showing population (blue spheres), known pathogenic (red spheres), early-onset (green spheres) and late-onset (orange spheres) epilepsy associated variants.

**Fig. S4.**
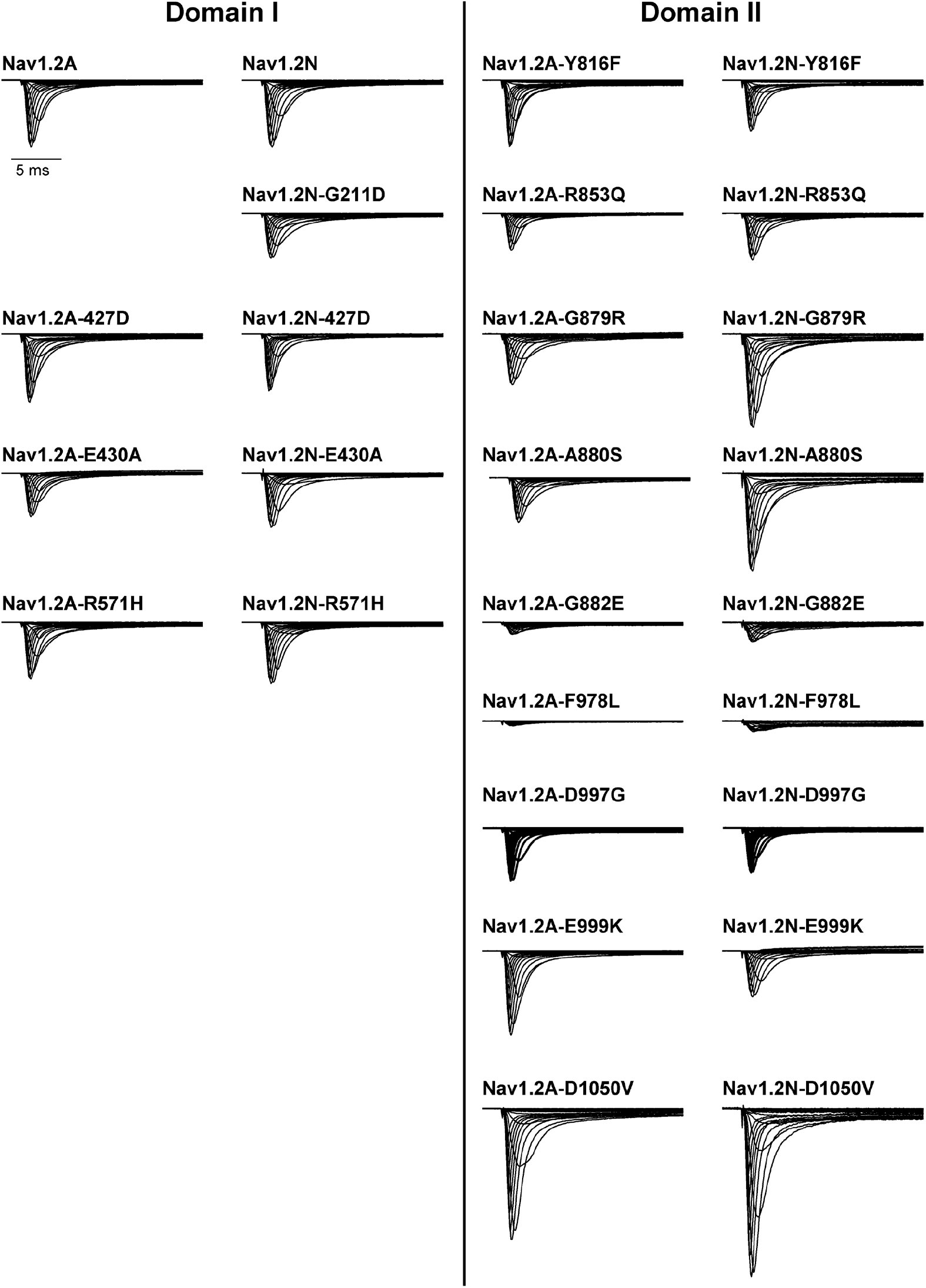
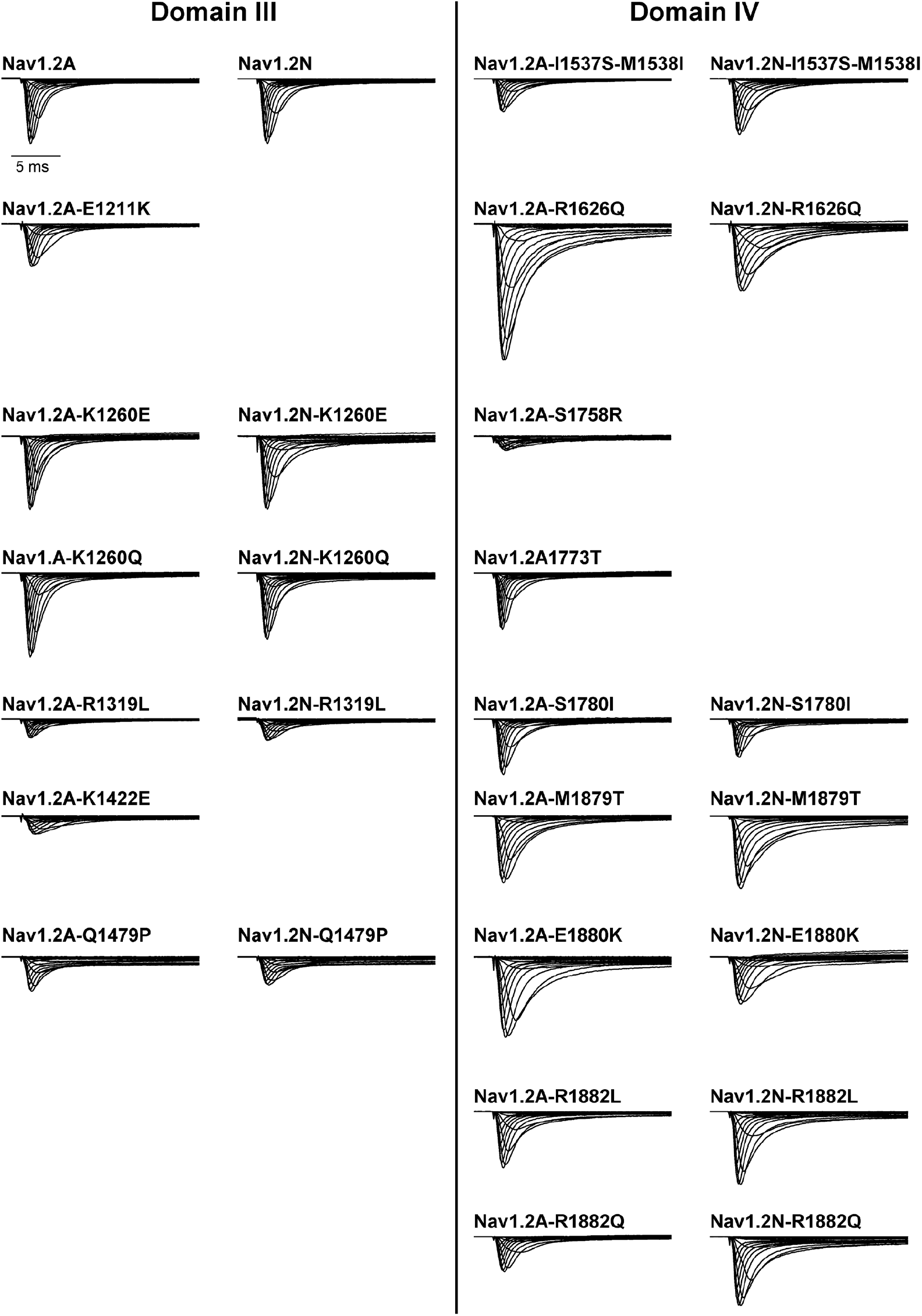
Average normalized whole-cell currents of Na_V_1.2 variants. Average normalized whole-cell sodium currents of Na_V_1.2 variants located in Domain I (left) and Domain II (right). Currents from the adult and neonatal splice isoform of each variant are shown side-by-side. All average traces were from 5 to 79 cells.

**Fig. S5.**
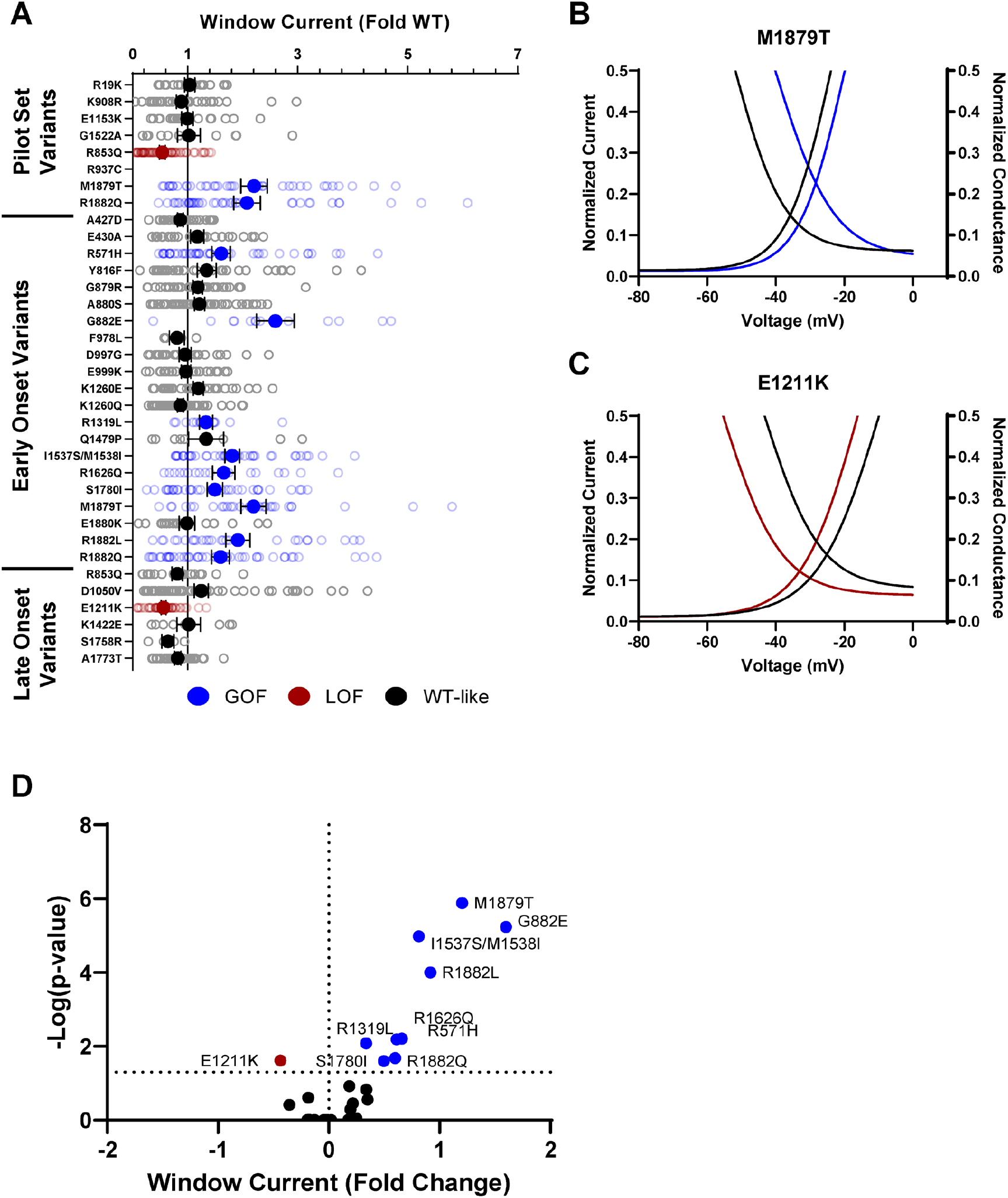
Na_V_1.2 variants affect window current. **(A)** Average deviation from WT Na_V_1.2 for the window current area. Boltzmann fit lines of representative variants showing **(B)** GoF; M1879T or **(C)** LoF; E1211K window current respective to WT. **(D)** Volcano plot highlighting variants significantly different from WT. Red symbols and lines denote loss-of-function and blue symbols and lines denote gain-of-function with p < 0.05 (n = 5-57).

**Fig. S6.**
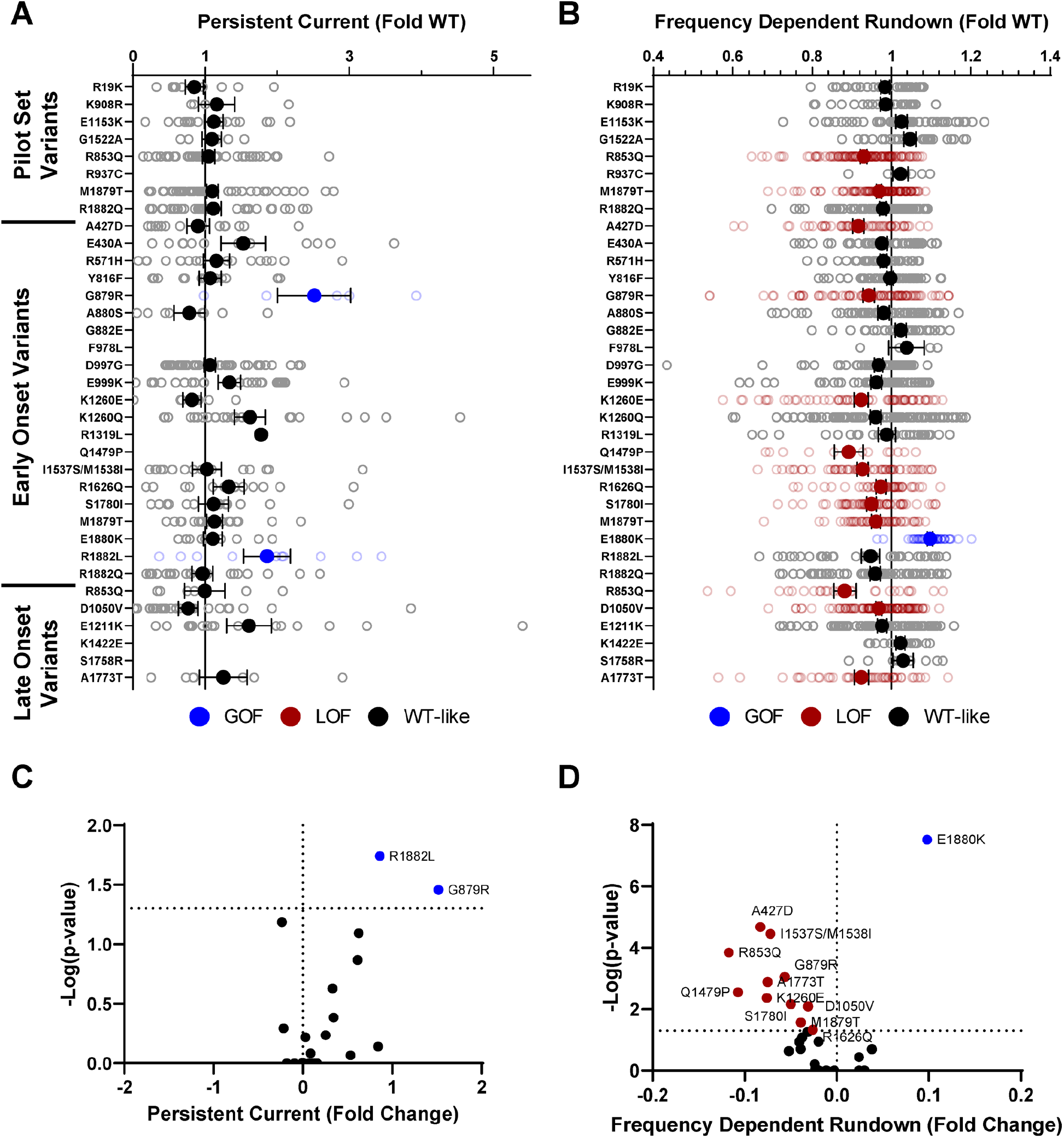
Na_V_1.2 variants affect channel inactivation. Average deviation from WT Na_V_1.2 for **(A)** persistent current and **(B)** frequency-dependent channel rundown at 20 Hz. Volcano plots highlighting variants significantly different from WT for **(C)** persistent current and **(D)** frequency-dependent rundown. Red symbols denote loss-of-function and blue symbols denote gain-of-function with p < 0.05 (n = 5-103).

**Fig. S7.**
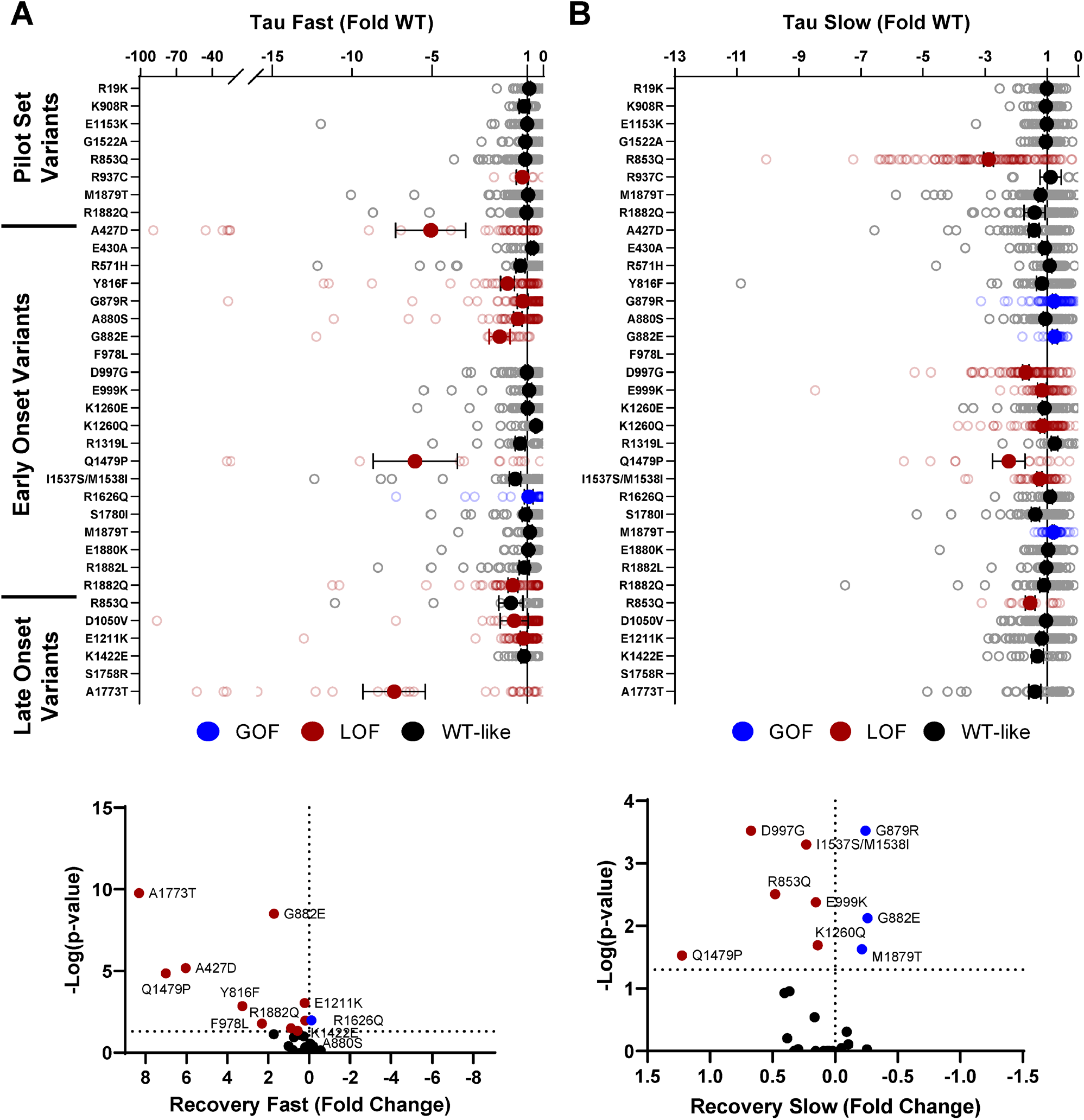
Na_V_1.2 variants affect recovery from inactivation. Average deviation from WT Na_V_1.2 for the **(A)** fast and **(B)** time constants of recovery from inactivation. Volcano plots highlighting variants significantly different from WT for **(C)** fast and **(D)** slow time constants of recovery from inactivation. Red symbols denote loss-of-function and blue symbols denote gain-of-function with p < 0.05 (n = 13-97).

**Fig. S8.**
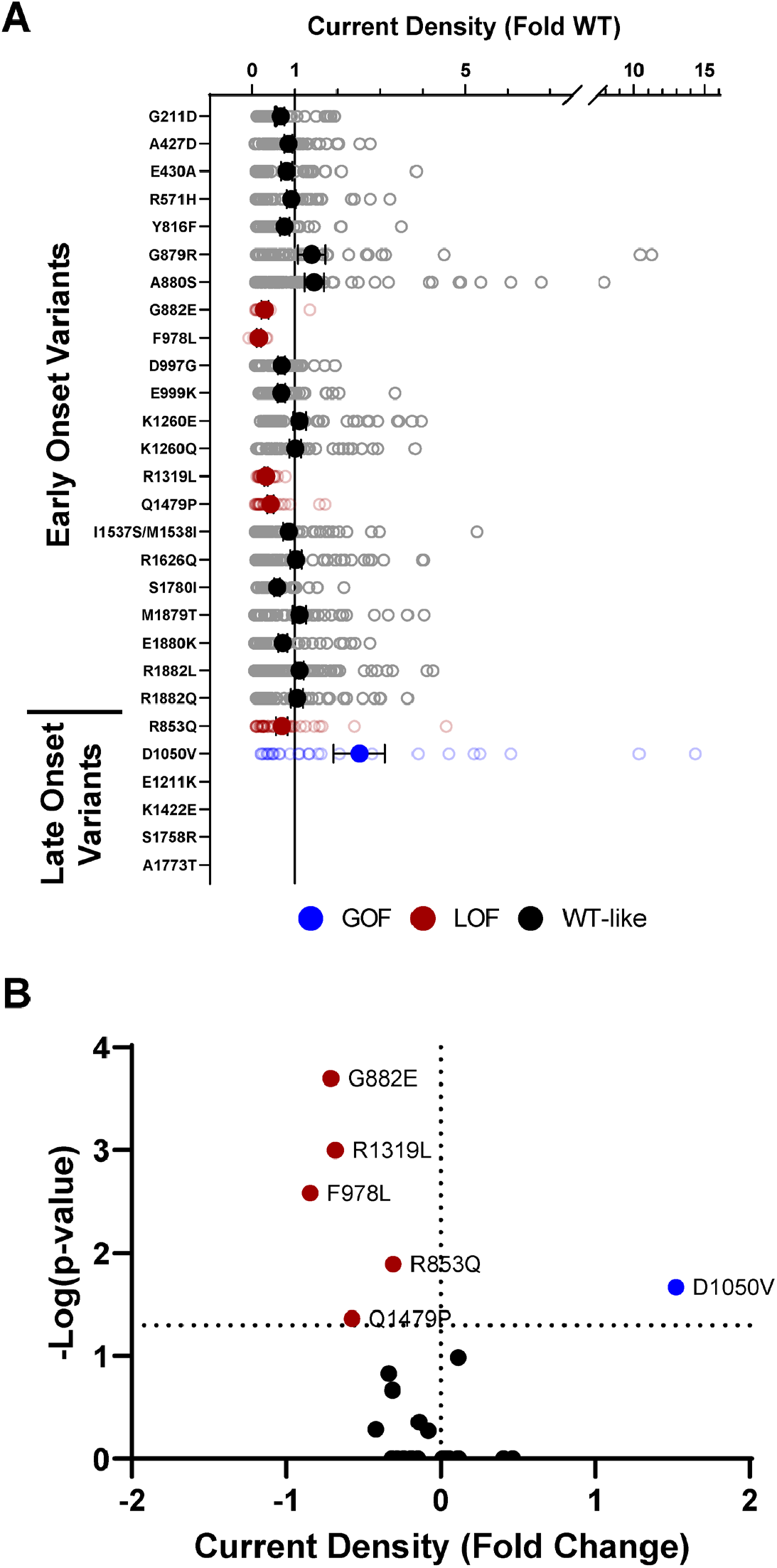
Disease-associated variants affect neonatal Na_V_1.2 whole-cell currents. **(A)** Average deviation of whole-cell sodium current density from neonatal WT Na_V_1.2 for epilepsy associated variants. **(B)** Volcano plot highlighting variants significantly different from WT. Red symbols denote loss-of-function and blue symbols denote gain-of-function with p < 0.05 (n = 9-81).

**Fig. S9.**
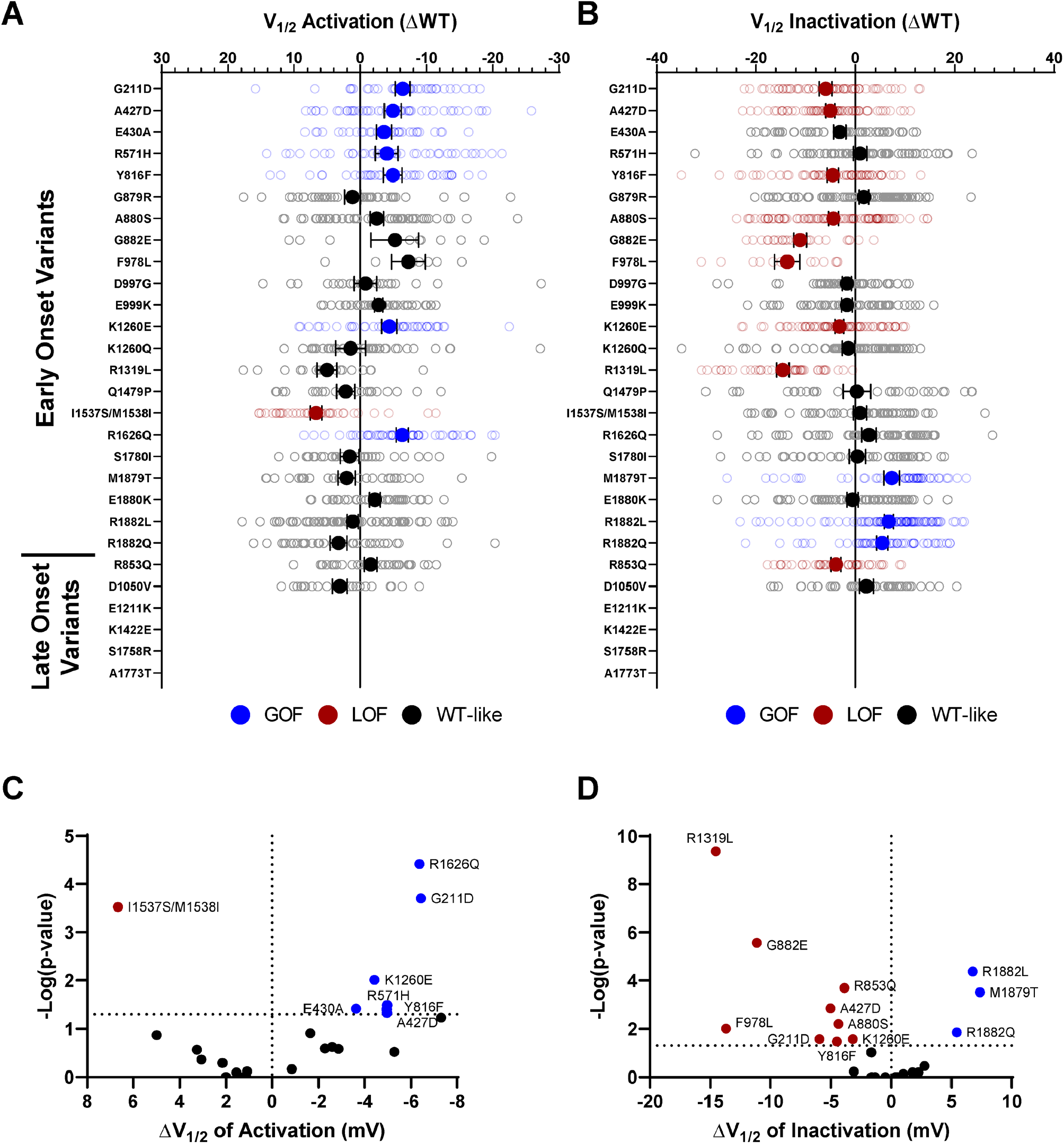
Disease-associated variants affect neonatal Na_V_1.2 voltage-dependence of activation and inactivation. Average deviation from neonatal WT Na_V_1.2 for V_1/2_ of **(A)** activation and **(B)** steady-state inactivation (in mV). Volcano plots highlighting variants significantly different from WT for voltage-dependence of **(C)** activation and **(D)** inactivation. Red symbols denote loss-of-function and blue symbols denote gain-of-function with p < 0.05 (n = 7-95).

**Fig. S10.**
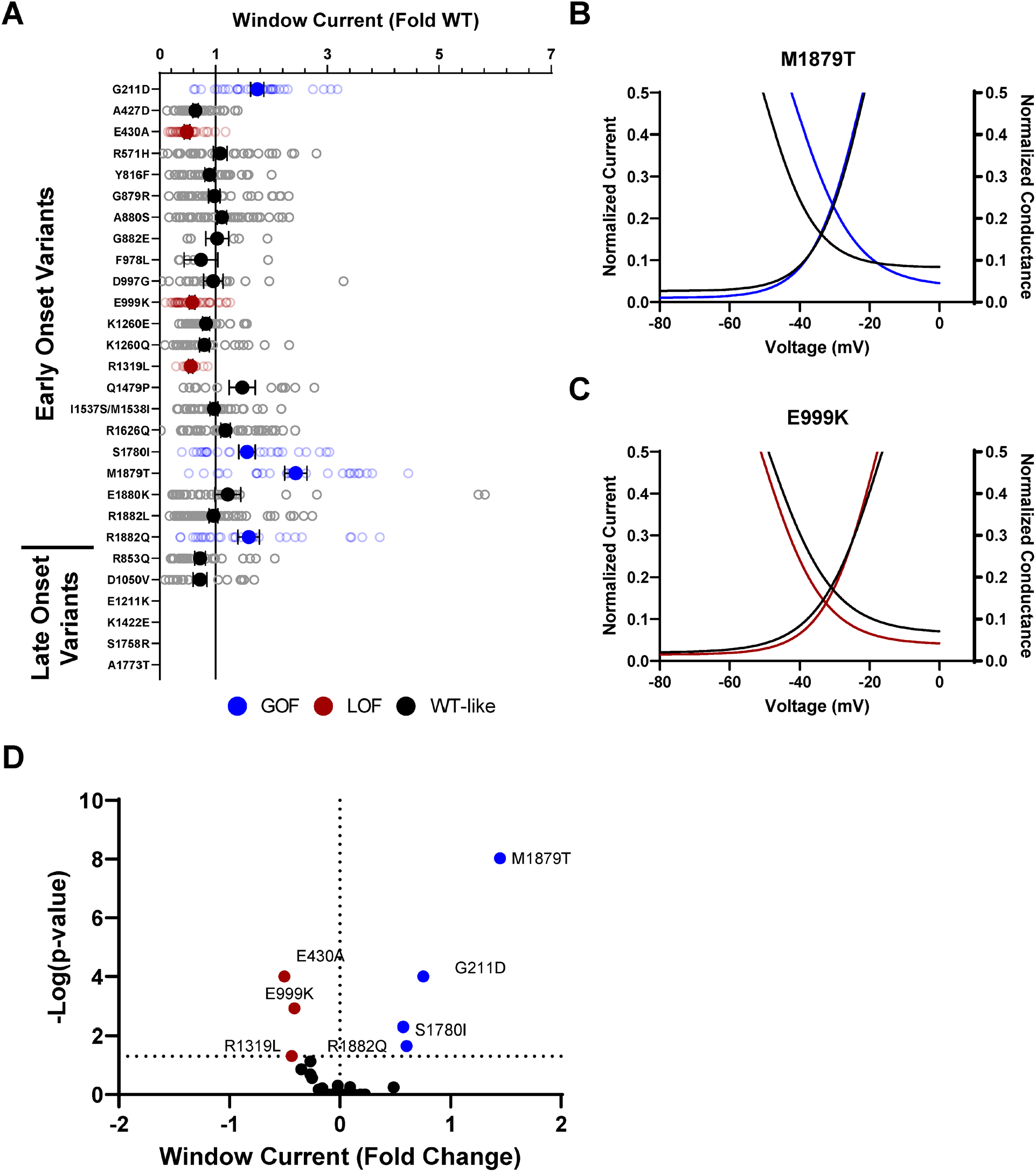
Disease-associated variants affect neonatal Na_V_1.2 affect window current. **(A)** Average deviation from WT Na_V_1.2 for the window current area. Boltzmann fit lines of representative variants showing **(B)** GoF; M1879T or **(C)** LoF; E999K window current respective to WT. **(D)** Volcano plot highlighting variants significantly different from WT. Red symbols and lines denote loss-of-function and blue symbols and lines denote gain-of-function with p < 0.05 (n = 5-59).

**Fig. S11.**
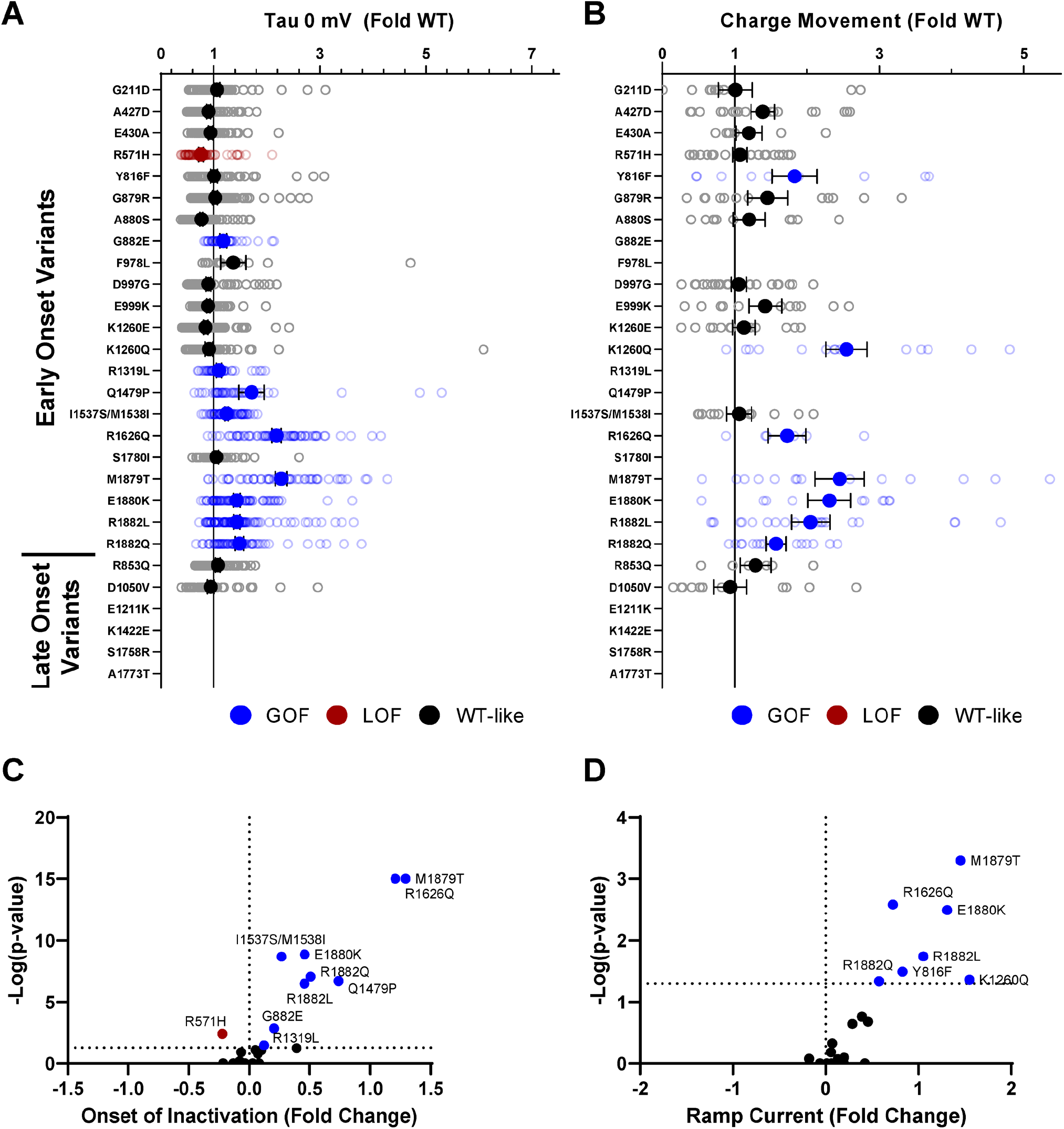
Disease-associated variants affect neonatal Na_V_1.2 affect inactivation time constants and ramp currents. Average deviation of **(A)** inactivation time constant (tau) and **(B)** ramp currents from neonatal WT Na_V_1.2 for epilepsy associated variants. Red symbols denote loss-of-function and blue symbols denote gain-of-function with p < 0.05 (n = 6-101).

**Fig. S12.**
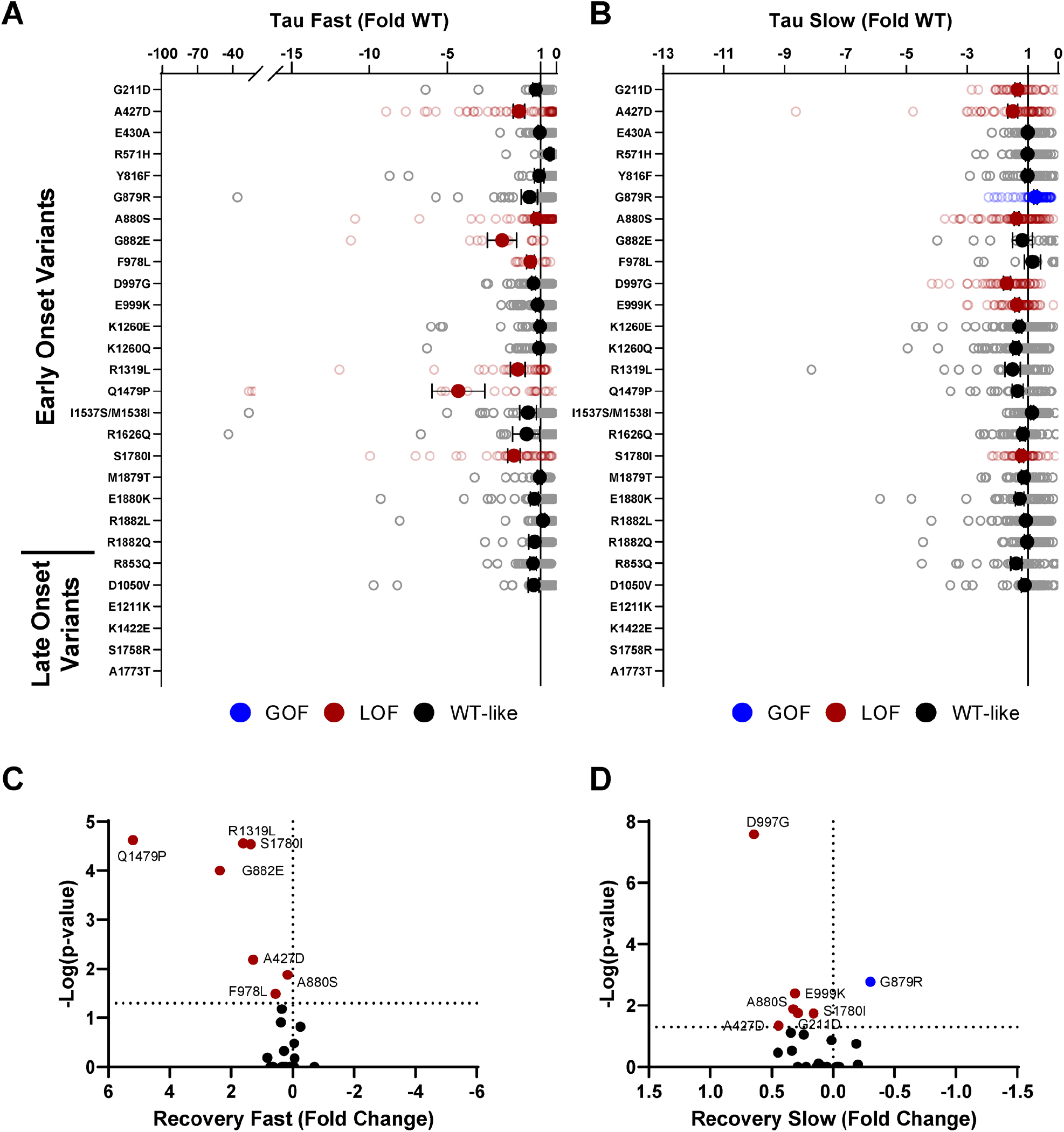
Disease-associated variants affect neonatal Na_V_1.2 affect recovery from inactivation. Average deviation from neonatal WT Na_V_1.2 for the **(A)** fast and **(B)** time constants of recovery from inactivation. Volcano plots highlighting variants significantly different from WT for **(C)** fast and **(D)** slow time constants of recovery from inactivation. Red symbols denote loss-of-function and blue symbols denote gain-of-function with p < 0.05 (n = 12-84).

**Fig. S13.**
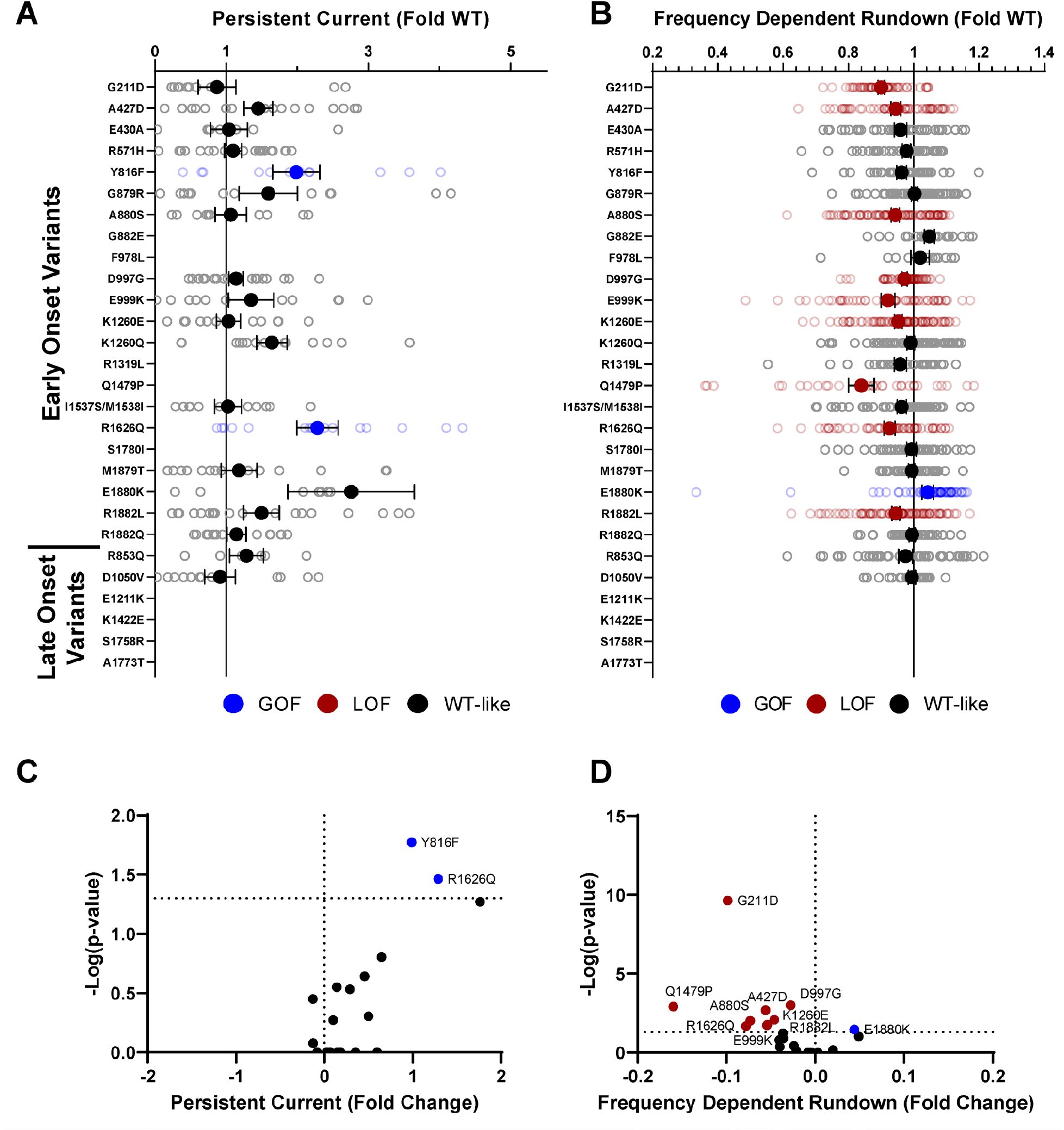
Disease-associated variants affect neonatal Na_V_1.2 affect persistent current and frequency-dependent channel rundown. Average deviation from neonatal WT Na_V_1.2 for **(A)** persistent sodium current and **(B)** frequency-dependent channel rundown at 20 Hz. Volcano plots highlighting variants significantly different from WT for **(C)** persistent current and **(D)** frequency-dependent rundown. Red symbols denote loss-of-function and blue symbols denote gain-of-function with p < 0.05 (n = 6-88).

**Fig. S14.**
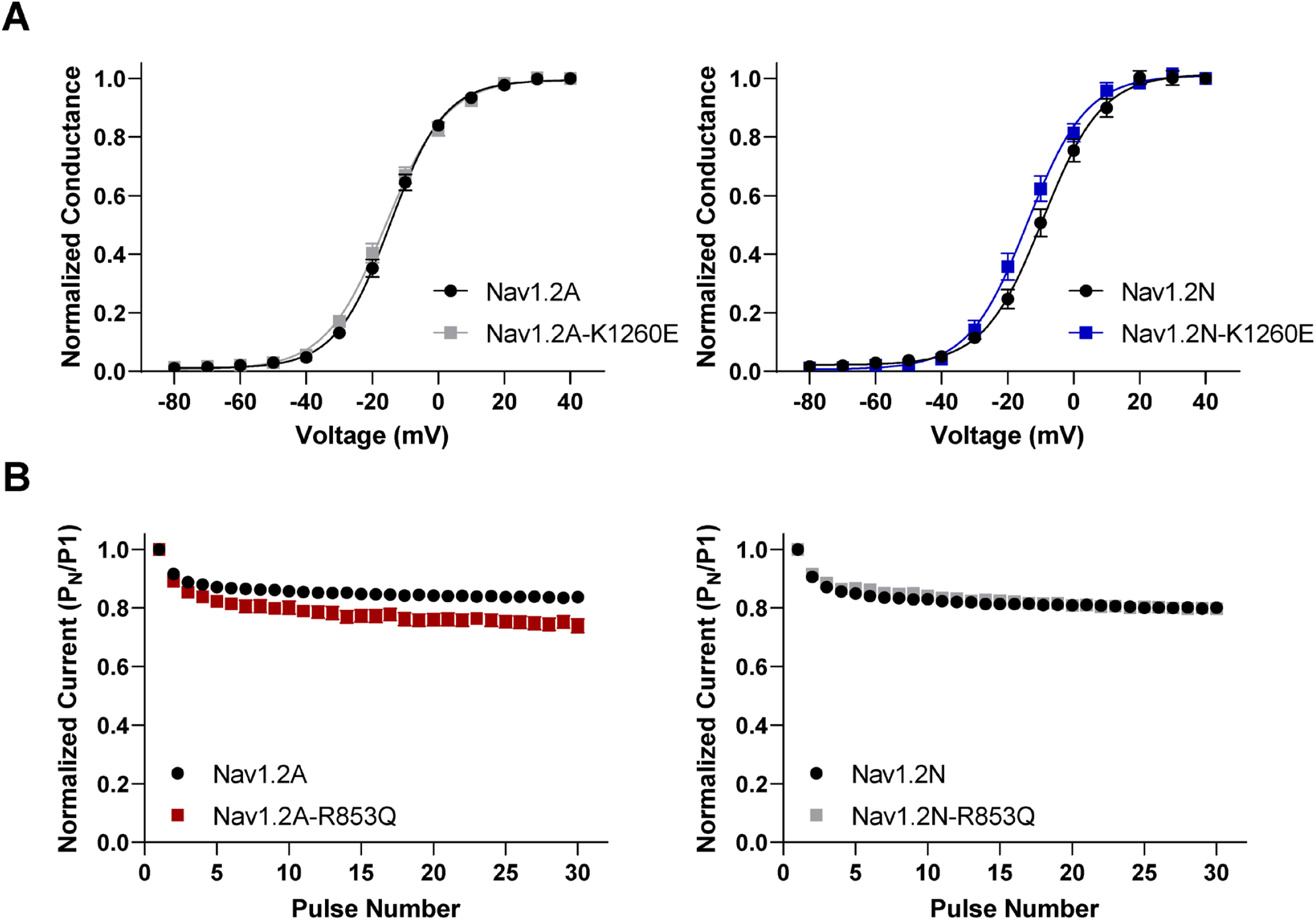
Na_V_1.2 variants exhibit splice isoform dependent functional properties. **(A)** Voltage dependence of activation of adult (left) and neonatal (right) splice isoforms of K1260E. **(B)**. Frequency-dependent channel rundown of adult (left) and neonatal (right) splice isoforms of R853Q. All data are expressed as mean ± SEM of 27 to 88 cells.

### Supplemental Tables

**Table S1.**
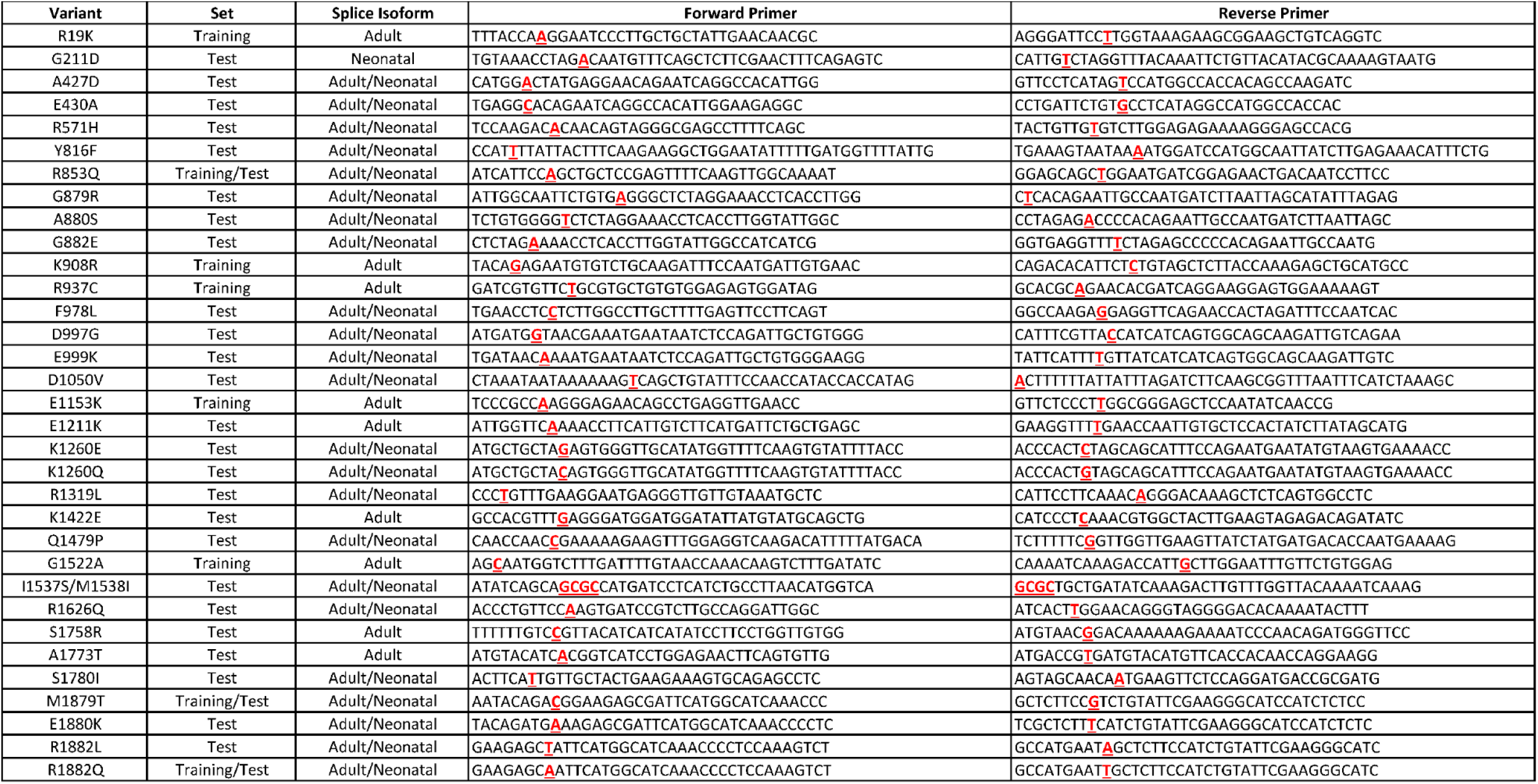
Mutagenic SCN2A primers

**Table S2.** Comparison of NaV1.2A biophysical properties

**Table S3.** Biophysical properties of NaV1.2A population and epilepsy associated variants

**Table S4.** Biophysical properties of NaV1.2A epilepsy associated variants

**Table S5.** Biophysical properties of NaV1.2N epilepsy associated variants

## Notes

### Competing Interest Statement

The authors have declared no competing interest.

